# Actomyosin pulsing rescues embryonic tissue folding from disruption by myosin fluctuations

**DOI:** 10.1101/2023.03.16.533016

**Authors:** Hongkang Zhu, Ben O’Shaughnessy

## Abstract

During early development, myosin II mechanically reshapes and folds embryo tissue. A much-studied example is ventral furrow formation in *Drosophila*, marking the onset of gastrulation. Furrowing is driven by contraction of actomyosin networks on apical cell surfaces, but how the myosin patterning encodes tissue shape is unclear, and elastic models failed to reproduce essential features of experimental cell contraction profiles. The myosin patterning exhibits substantial cell-to-cell fluctuations with pulsatile time-dependence, a striking but unexplained feature of morphogenesis in many organisms. Here, using biophysical modeling we find viscous forces offer the principle resistance to actomyosin-driven apical constriction. In consequence, tissue shape is encoded in the direction-dependent curvature of the myosin patterning which orients an anterior-posterior furrow. Tissue contraction is highly sensitive to cell-to-cell myosin fluctuations, explaining furrowing failure in genetically perturbed embryos whose fluctuations are temporally persistent. In wild-type embryos, this catastrophic outcome is averted by pulsatile myosin time-dependence, a time-averaging effect that rescues furrowing. This low pass filter mechanism may underlie the usage of actomyosin pulsing in diverse morphogenetic processes across many organisms.

## Introduction

Morphogenesis in the early embryo involves sculpting of tissue into specific shapes that are the precursors of organs and other specialized structures^1,2^. Orchestrated by biochemical signaling, mechanically coupled cells exert actomyosin contractile forces in supracellular networks to reshape tissue. The patterning of activated myosin II levels across cells determines the evolution of tissue shape, together with the elastic and viscous forces that oppose reshaping.

A fundamental challenge is to understand how a given myosin patterning leads to a certain tissue shape response. Myosin patterning is particularly well-characterized for ventral furrow formation (VFF) in the fruit fly *Drosophila*, a model organism for development studies. Commencing ∼ 3 hours after fertilization, VFF is the earliest major morphogenetic event^3,4^ and marks the onset of gastrulation, in which the embryo evolves from an ellipsoidally-shaped epithelial sheet of cells^5,6^ into a multilayered organization with the three germ layers: the ectoderm, mesoderm and endoderm. VFF, the first stage of mesoderm invagination, is driven by actomyosin networks on the apical surfaces of individual ventral cells, mechanically connected by intercellular adherens junctions^7^. Contraction of the networks constricts apical cell surfaces, an asymmetric deformation that creates tissue curvature and furrowing^8-11^ (Fig. 1A, B). The amount of apical medial myosin varies from cell to cell, in a ∼ 10-15 cell wide bell-shaped profile in the ventral-lateral direction^12-14^, extending ∼ 30 cells along the anterior-posterior axis with constant amplitude^7,14,15^. Thus, the myosin profile driving VFF has curvature in one direction, but no curvature in the other. Over ∼ 7 min the profile ramps up with little shape change^11,14^.

**Figure 1.**
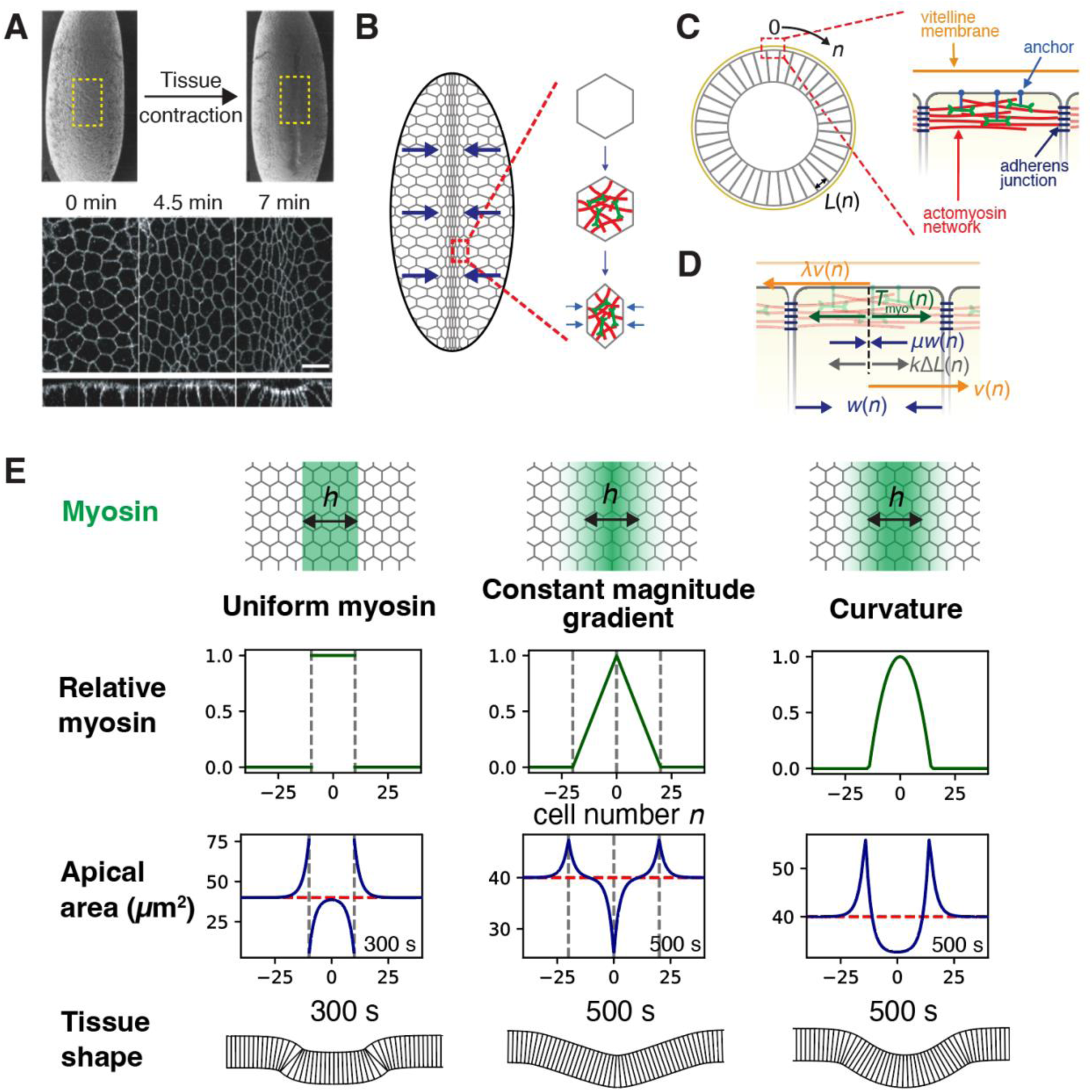
Curvature of the myosin envelope is required for tissue contraction and ventral furrowing in *Drosophila*. (A) *Drosophila* embryo before and after ventral furrow formation (top, electron micrographs adapted with permission from ref.^6^). Apical and transverse views of the presumptive mesoderm tissue at the indicated times relative to the onset of ventral furrow formation, corresponding approximately to the yellow boxes (bottom, fluorescence micrographs of labeled cell membranes, adapted with permission from ref.^10^; scale bar 10 μm.) (B) Cells assemble apical actomyosin networks whose contraction drives ventral furrowing, schematic. (C), (D) Mathematical model of *Drosophila* ventral furrow formation. Model of furrow formation. The *n*^th^ cell from the midline has width *L*(*n*), velocity *ν*(*n*) and contraction rate *w*(*n*). (C) Embryo cross-section (not to scale). Apical actomyosin networks are anchored to the apical membrane. The vitelline membrane is close to the apical cell surfaces. Adherens junctions connect neighboring cells. (D) Forces in the model. Actomyosin contractile tension *T*_myo_(*n*), internal viscous stress *μw*(*n*), external frictional drag from vitelline membrane *λν*(*n*), internal elastic stress *k L*(*n*). (E) Three hypothetical myosin profiles (top two rows) and the model-predicted cell area profiles (third row) and tissue shapes (bottom row) at the indicated times. Profiles are versus cell number *n*, the distance in cells from the ventral midline (Figs. 1(B), (C)). Initial cell areas, 40 μm^2^ (dashed red lines). Myosin profiles were maintained constant in time. Without curvature, substantial apical cell contraction or expansion is confined to the singular points of the profile (gray dashed lines). Only the profile with curvature apically contracts a broad band of cells, generating a smoothly shaped furrow with extended curvature. Model parameters as in Table 1.

To explain how this myosin patterning over the ventral embryo surface leads to furrowing, mathematical models using continuum or vertex model frameworks invoked elastic forces that resist the actomyosin-driven cell compression^11,13,16-18^. These elasticity-based models predict collective cell contraction and furrowing, but have serious shortcomings. First, the experimentally reported elastic constant for apical cell constriction is ∼ 7 pN μm^−119^, far too small to resist the myosin contractile forces of ∼ 1-1.5 nN exerted by a *Drosophila* apical network^20^. Second, in wild type embryos the myosin profile during VFF has no curvature in the anterior-posterior direction, and cells do not contract in that direction^7^. Elastic models fail to capture this behavior^11^, since they predict collective cell contraction even for a flat myosin profile. To curtail anterior-posterior contraction, some elastic models used unphysiological fixed cell boundary conditions^16^. Third, elastic models^11,18^ fail to reproduce the observed apical cell area profile in the presumptive mesoderm during VFF, in particular the ∼ 3 cell-wide^12^ cell expansion zones at the furrow periphery. Some elastic models did reproduce these expansion zones, but were forced to take adhoc measures without experimental basis, including fixed cell boundary conditions^16^, high elasticity zones near the midline^13^, or the use of myosin profiles^17^ inconsistent with experiment. Fourth, purely elastic models lack memory effects^16^. Memory is evident from experiments, e.g. apical cell areas lag myosin levels^21^, and partial recovery of cell areas required ∼ 2 min following treatment with an acute myosin-inhibiting drug during VFF^8^.

**Table 1.** Model parameter values.

| Symbol | Meaning | Value (mean ± standard error) | Legend |
| --- | --- | --- | --- |
| $\xi$ | Damping length | $2.5 \pm 1.2$ | (A) |
| $w^*$ | Characteristic contraction rate | $42 \pm 6 \text{ nm s}^{-1}$ | (B) |
| $\mu$ | Internal viscosity | $36 \pm 5 \text{ pN s nm}^{-1}$ | (C) |
| $\lambda$ | External drag coefficient | $6.3 \pm 6.1 \text{ pN s nm}^{-1}$ | (D) |

(C) From $\mu = T_{\text{myo}}^*/w^*$ , using the experimental myosin tension $T_{\text{myo}}^* = 1.5 \text{ nN}$ during dorsal closure<sup>20</sup>.
(D) From $\lambda = T_{\text{myo}}^*/(\xi^2 w^*)$ .

The myosin pattern that encodes a mechanical tissue response during VFF is remarkably robust to noise. Generally, protein levels in cells have significant fluctuations due to stochastic upstream gene expression, signaling cascades and biophysical processes^22-24^, and robustness to such effects as well as to genetic and environmental fluctuations is a necessary feature of robust development^24,25^. In the case of *Drosophila* VFF, the myosin profile has a smooth envelope but myosin levels vary stochastically by ∼ 45% about the envelope from cell to cell^12,14^. The fluctuations are both spatial and temporal: myosin in individual cells has pulsatile time-dependence, with sharp stochastic upward and downward variations over ∼ 60-90 sec superposed on a gradual ramp-up over ∼ 7 min^10,14^. Indeed, actomyosin pulsing and contraction pulses are a common feature of morphogenesis in many organisms^26-28^, including germband extension^29,30^ and dorsal closure^31-34^ in *Drosophila*, internalization of endoderm precursors in *C. elegans*^35^, and neurulation in *Xenopus*^36^. Pulsing is also a feature of actomyosin-driven polarization of individual cells^37^.

In summary, it is unknown how the potentially disruptive effects of myosin fluctuations are controlled during *Drosophila* VFF, and the function, if any, of pulsatile myosin time-dependence remains a mystery despite its prevalence. Moreover, significant evidence suggests dissipative effects beyond a purely elastic response must be invoked to explain the evolution of tissue shape during VFF. Here, we conclude that extended apical cell contraction is resisted primarily by viscous drag, with small elastic contributions. A biophysical model of *Drosophila* VFF shows that, in consequence, cell contraction and tissue furrowing are encoded in the anisotropic curvature of the myosin patterning, which orients an anterior-posterior furrow. This dependence on myosin profile curvature explains the absence of apical cell contraction in the anterior-posterior direction, and furrowing failure in Spn27A depleted embryos^14^ whose myosin profiles are flat. The model accurately reproduces the measured time-dependence of furrow depth and the experimental apical area profiles, including expansion zones which emerge naturally. We find that tissue contraction is highly sensitive to cell-to-cell myosin fluctuations: above a threshold fluctuation furrowing is disrupted, explaining furrowing failure in embryos with temporally persistent myosin fluctuations, either with depleted RhoGAP^38^ or in myosin phosphomutants^39^. In wild-type embryos, this catastrophic behavior is averted by pulsing of activated myosin: pulsing time averages the myosin fluctuations, suppressing the net fluctuation and rescuing furrowing by a low pass filter mechanism. We suggest this mechanism explains the usage of pulsatile myosin dynamics in diverse morphogenetic processes across many organisms^29-36,40-42^.

## Results

### Model

Following several rounds of division of the *Drosophila* zygote nucleus, ∼ 6000 nuclei migrate to the plasma membrane to yield the syncytial blastoderm^43^. Cellularization begins ∼ 2 hours after fertilization and is complete within ∼ 1 hour (Fig. 1A), when VFF onsets. VFF is the first major morphogenetic event, when contraction of the presumptive mesoderm along the ventral-lateral axis generates a ventral furrow over ∼ 7 min (Fig. 1A). During the subsequent ∼ 8 min the presumptive mesoderm invaginates into the embryo, ultimately developing into muscles, heart, fat bodies and other structures^44^.

We built a mathematical model of VFF. Contraction and furrowing of the epithelial cell sheet is driven by contractile actomyosin networks on the apical surfaces of ventral cells, with variable levels of activated myosin (Fig. 1B, C). In the activated myosin region that extends ∼ 30 cells along the anterior-posterior axis, the apical areas, contraction rates, velocities and myosin levels of cells are statistically independent of location in the anterior-posterior direction^6,7,14^. Thus, for simplicity we consider variations of cell properties along the ventral-lateral axis only. The force balance on the *n*^th^ cell from the ventral midline describes a balance of actomyosin, viscous and elastic forces (Fig. 1D). Taking the continuous limit gives

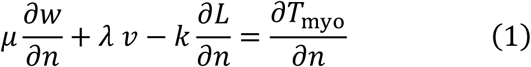

where *ν*(*n*), *w*(*n*) and *L*(*n*) are, respectively, the cell velocity, contraction rate and width, and *T*_myo_(*n*) is the actomyosin stress, assumed proportional to the amount of myosin whose profile has a bell-shaped envelope in the ventral-lateral direction, superposed on which are substantial cell-to-cell variations^12,14^. The internal viscosity *μ* reflects network remodeling and other dissipative processes accompanying cell contraction, and 2*λ* is the external drag coefficient opposing translational cell motion, likely dominated by interactions with the nearby vitelline membrane^19,45^. The elastic constant *k* measures elastic stresses resisting apical area change, experimentally reported to be *k* = 7 pN μm^−119^. Given cell widths ∼ 7 μm^10^, even a 100% strain in this very particular deformation mode would provoke an elastic stress ∼ 50 pN, far smaller than the reported apical myosin contractile stresses of ∼ 1.5 nN (see Materials and Methods and Discussion for further discussion of this point). Thus we discard the elastic term. Taking the derivative, eq. (1) then gives

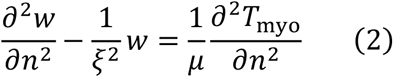

where *ξ* = (*μ*/*λ*)^1/2^ emerges as a damping length, whose significance is that external drag dominates scales larger than *ξ*.

All model parameters were determined from experiment (see Materials and Methods and Fig. S1). Fitting model-predicted apical area profiles to experiment yielded *ξ* = 2.5 ± 1.2 cells, *λ*∼6 pN s nm^−1^, and *μ*∼36 pN s nm^−1^ (Table 1). In the following sections, using experimental myosin profiles *T*_myo_(*n*) as input to Eq. (2) we solve for the contraction rate profile *w*(*n*) and hence cell width and area profiles *L*(*n*), *A*(*n*). Contraction of apical cell surfaces generates curvature and furrows the ventral surface (Fig. 1A); from the predicted apical area profile, we calculate the furrowed tissue shape (see Materials and Methods and Fig. S2). We also developed a fully 2D analysis explicitly treating variations in the anterior-posterior direction, which confirmed our conclusions from the above model (Materials and Methods).

### Tissue contraction and furrowing require curvature of the myosin envelope

The myosin patterning that drives VFF has a bell-shaped envelope in the ventral-lateral direction whose amplitude grows in time^12-14^. Is this particular envelope shape functionally significant? What are the requirements on the envelope for successful spatially extended tissue contraction and folding?

To gain insight, we first considered three simplified hypothetical myosin envelope shapes of width 20 cells in the ventral-lateral direction, fixed in time: a top-hat profile with constant amplitude in the central active region, a triangular profile with constant magnitude gradient in this region, and a parabolic profile which unlike the other two profiles has curvature (Fig. 1E). For each profile *T*_myo_(*n*) we solved Eq. (2) to obtain the cell contraction rates and apical areas versus cell number *n*, and from these we calculated the furrow shapes (Materials and Methods) (Fig. 1E). The top-hat and triangular myosin profiles have high amplitudes in the central active region, yet generated almost no cell contraction in this region except at isolated locations. Both result in badly misshapen furrows. Only the parabolic profile smoothly and collectively contracted the cells in the active region, generating a smoothly rounded furrow similar to that observed in embryos (cf. Figs 1E and 1A). Two cell expansion zones peripheral to the contracted active region are also generated, reproducing a distinctive feature of experimental apical cell area profiles during VFF (see later sections). Indeed, the expansion zones are required to smoothly join the furrow to the unfurrowed region (Fig. 1E).

The physical origin of the curvature requirement is that large scales (> *ξ*) are dominated by drag forces, likely from motion relative to the vitelline membrane. Since apical contraction of a row of cells requires their translational at different velocities (uniform translation of a group of cells has no effect on cell width), the actomyosin force must have gradient. Since the force is proportional to the amount of myosin, the myosin profile requires curvarture.

Thus, curvature in the myosin patterning is required for extended tissue contraction and well-shaped furrows. Activation of myosin is not in itself sufficient, nor is a myosin gradient.

### Myosin patterning with anisotropic curvature generates a ventral furrow

These results suggest that the curvature of the bell-shaped myosin profile in *Drosophila* embryos is the key property driving VFF. Indeed, solving Eq. (2) using the myosin envelope measured in wild type embryos during VFF in ref.^12^, the model quantiatively reproduced the experimental profile of apical areas of cells and the furrow shape, with contracted cells near the ventral midline, expanded cells in the wings, and a rounded furrow^12^ (Fig. 2A, left). Using the time-dependence of the experimentally measured myosin envelope as it ramps up over ∼ 500 sec^14^, the model accurately reproduced the measured increasing furrow depth over time^10^ (Fig. 2B) (Supplementary Information).

**Figure 2.**
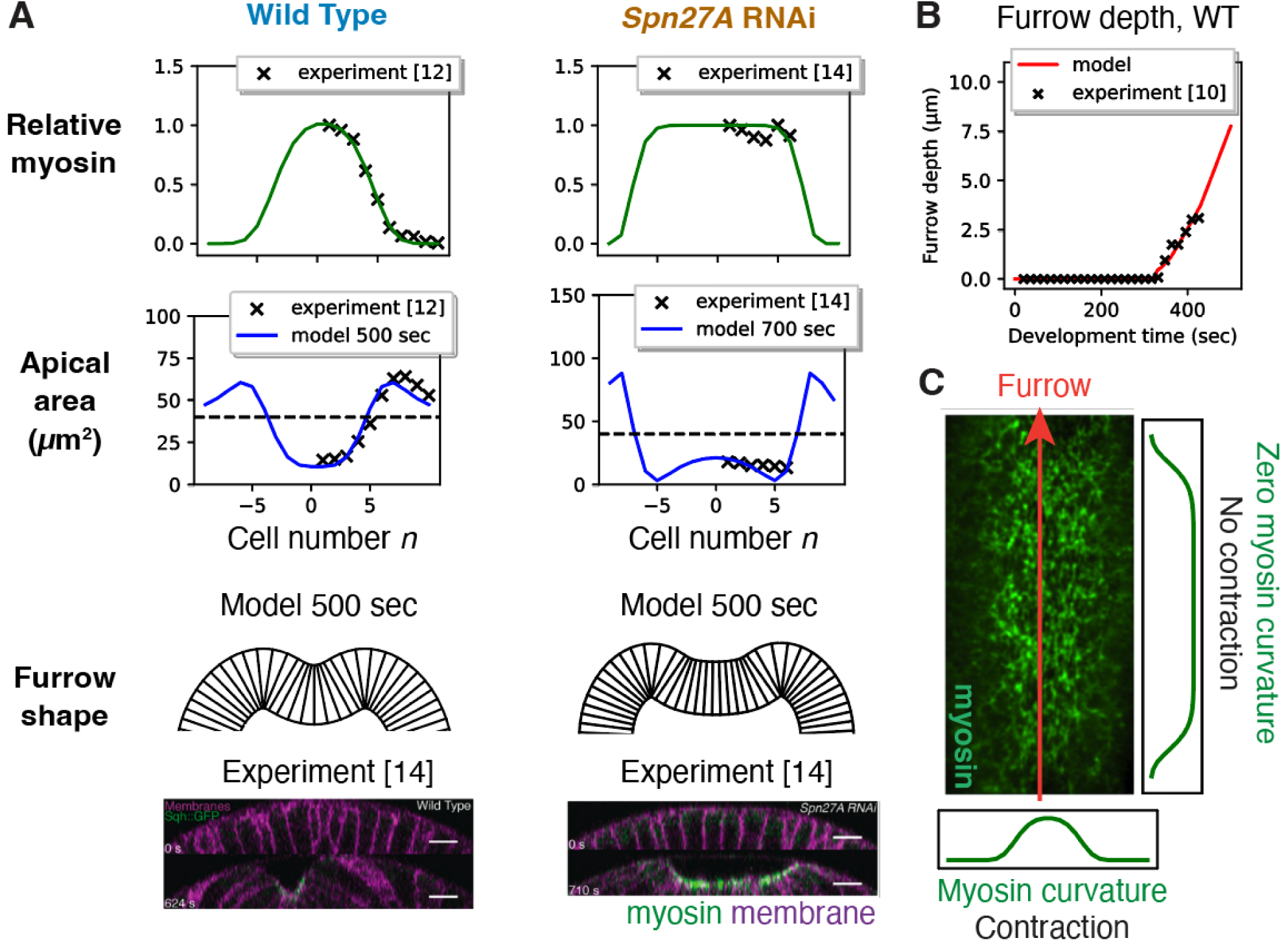
Anisotropic curvature of the myosin profile sculpts an oriented furrow during ventral furrow formation. (A) Using the experimental myosin profiles as input, the model reproduces the experimental apical area profile and furrow shape in wild-type and *Spn27A* RNAi embryos. Experimentally measured ventral myosin and cell apical area profiles of wild-type (from ref.^12^) and *Spn27A* RNAi (from ref.^14^) embryos (top 2 rows). Green curves: best fit myosin profiles, input to model. Blue curves: model predicted apical area profiles after 500 sec (wild-type) and 700 sec (*Spn27A* RNAi). Dashed lines: initial apical areas (40 μm^2^). The predicted wild-type apical area profile has a broad band of contracting cells, giving a furrow with extended curvature (bottom), whereas for *Spn27A* RNAi a paucity of strongly contracted cells near the midline generates an abnormally expanded and flat furrow (bottom). Predicted furrow shapes are close to the experimental shapes (fluorescence micrographs adapted with permission from ref.^12^). Model parameters as in Table 1. (B) Furrow depth versus time predicted by the model (red curve) using as input the time-dependent ventral myosin profile measured in ref.^12,14^. Experimental measurements of furrow depths (crosses) from ref.^10^. (C) The experimental myosin envelope has curvature in the ventral-lateral direction, but is flat along the anterior-posterior axis (fluorescence micrograph adapted with permission from ref.^15^). This anisotropic curvature produces ventral-lateral contraction only, sculpting the tissue into a furrow oriented along the anterior-posterior axis.

The curvature requirement explains another important experimental observation: while apical surfaces contract in the ventral-lateral direction, no contraction occurs in the anterior-posterior direction^7^. This is as expected, since the myosin envelope has zero curvature in the latter direction^7,14^. To explicitly demonstrate this, we extended our analysis to a fully 2D continuum model incorporating internal and external dissipative forces and accounting for the full myosin envelope extending ∼30 cells along the anterior-posterior axis with almost constant amplitude^6,7,14^ (see Materials and Methods and Supplementary Information). Indeed, the predicted cell area profiles (Fig. S3) show no anterior-posterior contraction, and profiles in the ventral-lateral direction confirm the results of Fig. 2A from the model considering variations in this direction only.

Thus, our model (Eqs. (2), (3)) incorporating dissipative forces explains the experimental cell apical area profiles. Elastic models cannot reproduce these profiles without invoking unphysiological fixed cell boundary conditions^11,13,16^. To illustrate this, we added a substantial elastic term to the model of Eq. (2); the lateral cell expansion zones then failed to stabilize, and in contradiction with experiment continued to grow in size throughout VFF (Fig. S4).

In summary, the anisotropic curvature^7,14,15^ of the myosin envelope sculpts a furrow with specific orientation: the direction with curvature sets the direction of contraction and furrowing (ventral-lateral), while the direction of zero curvature defines the direction of zero contraction and furrow orientation (anterior-posterior) (Fig. 2C).

### Spn27A depletion leads to abnormal furrows because the myosin envelope lacks curvature

If furrowing requires curvature in the myosin patterning, we would expect blocked or abnormal furrowing if the curvature is suppressed. Such a myosin profile lacking curvature is realized in *Spn27A* RNAi embryos. Spn27A is a negative regulator of ventral cell fate at the edges of the myosin-enriched active zone^46^, so Spn27A depletion leads to a flattened ∼ 13 cell wide myosin envelope in the vental-lateral direction that drops sharply over 1-2 cells^14^ (Fig. 2A, right), close to the hypothetical top-hat shape considered above (Fig. 1E).

Consistent with the curvature requirement, in *Spn27A* RNAi experiments furrowing is abnormally expanded and flat (Fig. 2A, right), and further mesoderm invagination is usually blocked^14^. Moreover, the entire active zone detaches from the vitelline membrane, possibly due to severe apical constriction of cells at the very high curvature active zone edges.

To model these experiments, as input we used a myosin envelope with the top hat-like shape of the *Spn27A* RNAi embryo envelope (Fig. 2A, right). We assumed a reduced drag coefficient in the detached region (cells -6 to +6) to account for the vitelline membrane being distant. A best fit of the predicted apical area profile and furrow shape to experiment was obtained for a drag reduction factor *β* = 0.5 (Fig. S5). Consistent with experiment^14^, the predicted furrow shape is much flatter and wider than wild type and has high curvature at the active zone edges (Fig. 2A, active zone marked green, bottom left). This shape is similar to the expanded, flat furrow shape predicted for the hypothetical top hat myosin envelope (Fig. 1E). Overall, Spn27A depletion studies support the proposal that myosin curvature is required for functional furrow formation.

### Persistent cell-to-cell myosin fluctuations disrupt tissue contraction and furrowing

Thus far we considered the smooth myosin envelope only. In reality, myosin levels fluctuate significantly from cell to cell about this envelope^12-14^, an inevitable consequence of multiple intrinsically stochastic upstream processes, including the expression of genes such as *fog* and *T48* across the presumptive mesoderm^47^, and endocytosis of the GPCRs Mist and Smog^48^. The net effect is that myosin levels fluctuate *ϵ*_fluc_ ≈ 45% about the envelope (Supplementary Information and Fig. S6A, B).

What is the impact of these fluctuations? We first considered a single fluctuation whose relative value is frozen in time. Solving our model with a hypothetical myosin profile consisting of the experimental envelope^12^ plus a single deficit fluctuation of representative relative magnitude *ϵ*_fluc_ = 45%, the persistent myosin deficit not only prevented contraction but actually expanded the cell ∼ 30% after 400 s, producing a local defect in furrow shape (Fig. 3A).

**Figure 3.**
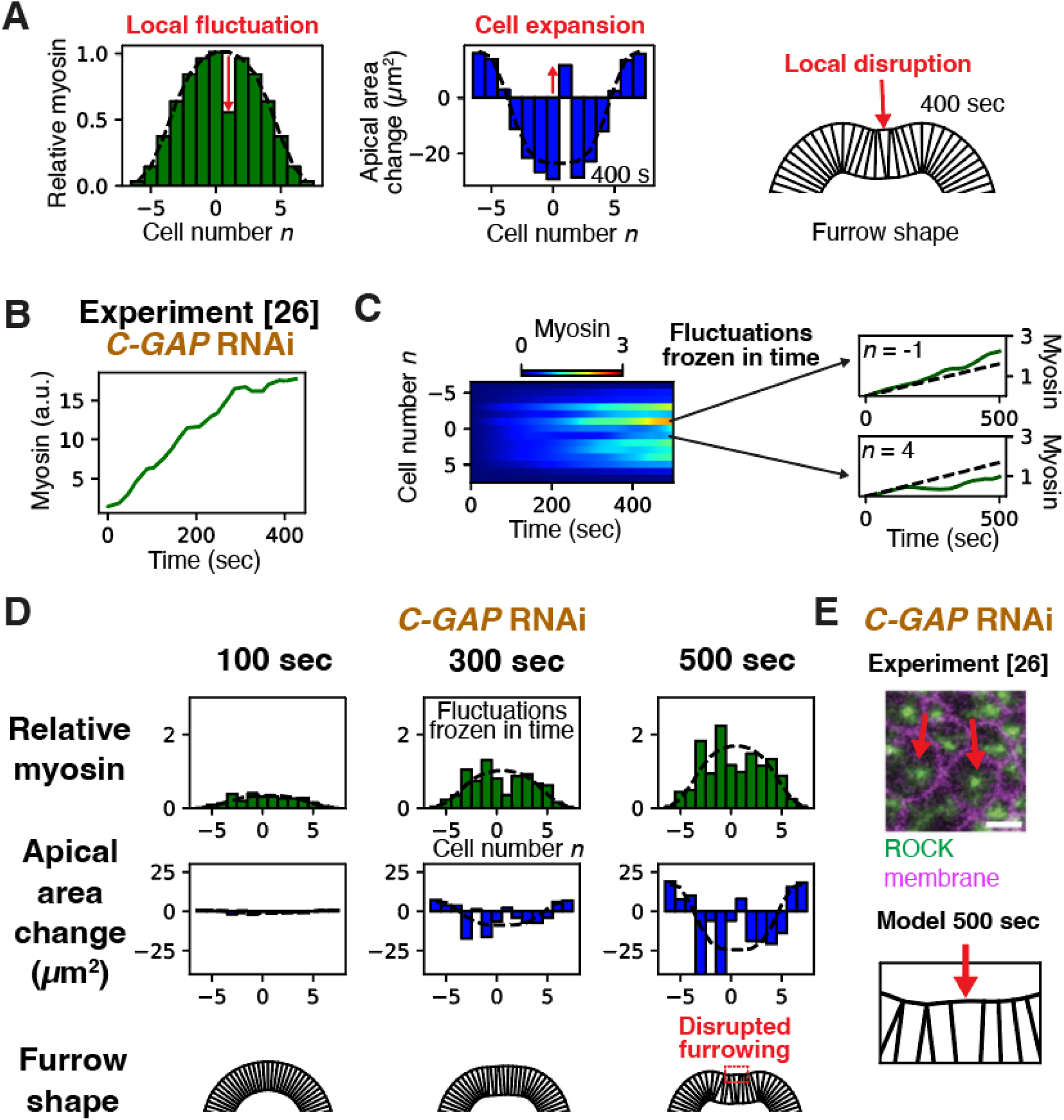
Cell-to-cell myosin fluctuations disrupt tissue contraction and furrowing in C-GAP depleted embryos. (A) Model-predicted apical area profile and tissue shape generated by a hypothetical myosin profile with a fluctuation at a single cell about the smooth envelope (dashed curve, experimental wild-type myosin envelope of Fig. 2A). The defecit fluctuation prevents apical constriction of that cell, with consequences for tissue shape (right). (B) Experimental myosin vs. time for a cell in a *C-GAP* RNAi embryo during VFF, adapted with permission from ref.^38^. Myosin ramps up at almost constant rate. (C) Myosin spatiotemporal profile used as input to model *C-GAP* RNAi embryos, closely mimicking experiment. Myosin in a given cell increases at almost constant rate. The rate of increase varies randomly from cell to cell. (D) Model results for *C-GAP* RNAi embryos, using the myosin profile of (C) as input. Relative cell-to-cell fluctuations of myosin are almost frozen in time, generating large cell area fluctuations. Homogeneous apical constriction over a central band of cells is lost, with some cells expanding. Furrowing is seriously disrupted. (E) Fluorescence micrograph of ventral cells in a *C-GAP* RNAi embryo during VFF, adapted with permission from ref.^38^ (top). Some cells fail to apically constrict (red arrows). The model reproduces apical constriction failure (bottom, blow-up of red box in (D); red arrow indicates an expanded cell).

This result suggests that when myosin fluctuations are frozen in time, apical constriction and furrowing are disrupted. This is consistent with experimental studies of embryos with depleted C-GAP, a Rho GTPase-activating protein (RhoGAP)^38^. In normal embryos, myosin pulses stochastically about the envelope due to cyclic activation and deactivation of the upstream RhoA by, respectively, the guanine nucleotide exchange factor RhoGEF2 and C-GAP^49^. Depleting C-GAP disrupts these RhoA dynamics, so in a given cell myosin ramps up in time but its amplitude relative to the envelope remains constant or changes slowly^38^ (Fig. 3B). Thus, in *C-GAP* RNAi embryos myosin fluctuations are persistent: the level in a given cell tends to remain above or below the envelope throughout VFF. In these embryos tissue contraction is inhomogeneous, with some ventral cells even expanding (Fig. 3E, arrows), and the invagination failure rate is high^38^. Myosin fluctuations are similarly persistent in phosphomimetic *sqh-EE or sqh-AE* mutant embryos with constitutively activated myosin, and in embryos with depleted myosin phosphatase. In the latter, cell apical area profiles are again highly heterogenous, while in phosphomimetic mutants invagination is delayed^39^.

To model *C-GAP* RNAi embryos, we solved Eq. (2) using spatiotemporal myosin patterning that mimicks the experimental patterning^38^. The bell-shaped envelope ramps up linearly in time, and the myosin level in cell *n* equals the local envelope value plus a relative fluctuation *ϵ*_fluc_(*n, t*) that changes slowly in time with statistics taken from experiment^38^ (Fig. 3B) (Materials and Methods and Supplementary Information). The random fluctuation factors *ϵ*_fluc_(*n, t*) can be positive or negative (surfeit or deficit) and are statistically independent among cells. Thus fluctuations are frozen in time, i.e. the myosin pattern has fixed shape as it ramps up (Fig. 3C, D). Model-predicted apical areas are highly inhomogeneous, with some expanding cells (Figs. 3D) as observed in *C-GAP* RNAi embryos^38^ (Fig. 3E). Predicted furrows show severe disruption (Fig. 3D), with large variations and a ∼ 15% furrow failure rate (Fig. S7A), consistent with invagination failure rates in *C-GAP* RNAi embryos^38^.

Thus, persistent cell-to-cell variations in myosin levels about the envelope disrupt collective cell contraction and furrowing. The origin of this sensitivity is that on scales less than the damping length *ξ* ∼ 2-3 cells, cell contraction/expansion is resisted by internal viscosity while external drag forces are weak. Hence a cell’s contraction rate depends on its myosin level relative to that of nearby cells, essentially the local envelope value, so a cell *n* with a myosin deficit *ϵ*_flug_ would expand at rate 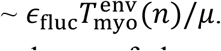. This local effect competes with the large scale curvature-driven contracting tendency of the envelope of width *h*∼8 cells, giving a contraction rate 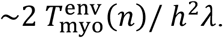. The local effect is greatest for fluctuations

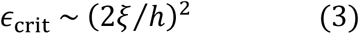

or greater. It follows that deficit fluctuations ∼ 25-55% are sufficient to expand a cell. Since measured fluctuations are of this order (Figs. 3B-D and S6A,B), cell-to-cell myosin variations pose a serious threat to extended apical cell constriction and furrowing.

### Pulsatile myosin fluctuations rescue furrowing by a low pass filter mechanism

How do wild-type embryos avoid the disrupted furrowing suffered by *C-GAP* RNAi embryos? A fundamental difference is that C-GAP depletion freezes the relative myosin fluctuations in time, whereas myosin in wild type cells executes dramatic pulsatile fluctuations, with a pulsing time ∼ 1 min^10^ and little correlation from cell to cell^10,21^ (Fig. 4A, B). Thus, the shape of wild-type myosin profiles is constantly changing, whereas the shape is persistent in *C-GAP* RNAi embryos (cf. Figs. 4C, 3D).

**Figure 4.**
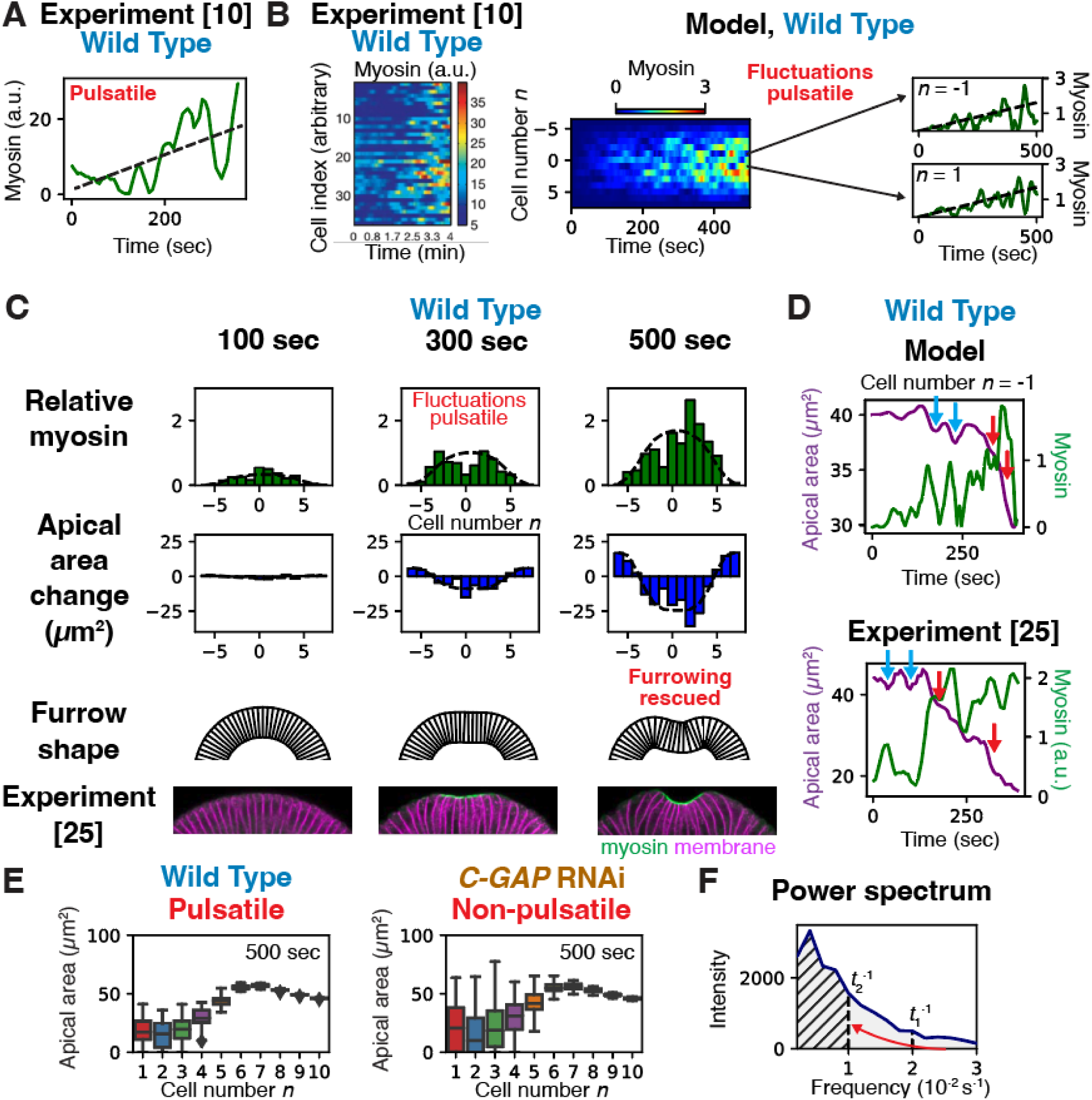
Pulsatile myosin dynamics rescue tissue furrowing from disruption by fluctuations in wild-type embryos. (A) Experimental apical myosin vs. time in a single of a wild-type embryo during VFF, showing large pulsatile variations about a monotonic ramp-up (dashed line). Adapted with permission from ref.^10^. (B) Experimental myosin levels of 40 ventral cells vs. time during VFF of a wild-type embryo, showing cells have statistically independent pulsatile dynamics. Adapated with permission from ref.^10^ (left). Kymograph of time-dependent myosin profile across 13 cells closest to the ventral midline, used as input to model VFF in wild-type embryos (right). The spatiotemporal profile quantitatively mimics experiment: cells have statistically independent pulsatile myosin fluctuations about th envelope (dashed lines). (C) Model results at the indicated times for the myosin profile of (B). With pulsatile time-dependence, all cells contract and furrowing is rescued (compare to Fig. 3D). (D) Myosin and apical area vs. time in a typical model cell (top) and experimental cell (bottom, adapted with permission from ref.^38^). Contraction pulses may be followed by episodes of expansion (blue arrows, “unratcheted”) or by further contraction only (red arrows, “ratcheted”). (E) Pulsatile myosin significantly lowers fluctuations in apical cell areas. Box-and-whisker plots compre model-predicted apical area profiles for wild-type and *C-GAP* RNAi embryos each with 50 myosin profiles of the type shown in (B) (wild-type, pulsatile) or Fig. 3C (*C-GAP* RNAi, non-pulsatile). (F) Pulsatile myosin suppresses fluctuations by a low pass filter mechanism. The plot shows the mean power spectrum of the pulsatile component of the myosin signal in the cell at the ventral midline (*n* = 1) used as input for the wild-type model, averaged over 50 realizations of the stochastic signal. The pulsatile component is the full signal minus the envelope value at *n* = 1. The total area (gray) represents the net myosin fluctuations about the envelope. At some time *t*_1_, the apical area profile depends on the time integral of the signal, a quantity whose power spectrum is suppressed for frequencies 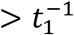. As time increases, more of the highest frequencies are filtered out (low pass filter). By *t*_2_ = 100 sec, the contributing myosin fluctuations are reduced to the hatched area. The reduced myosin signal fluctuations lower the apical cell area fluctuations and suppress disruption to furrowing.

We now show that it is indeed the pulsatile myosin time-dependence that rescues normal embryos from the disruptive effects of spatial myosin fluctuations^26-28^. We applied our model to wild-type embryos, repeating the *C-GAP* RNAi analysis but now the relative fluctuation of myosin about the envelope value, *ϵ*_fluc_(*n, t*), varies substantially in time, with a ∼ 1 min correlation yielding a pulsatile signal closely mimicking experiment^10,21,38^ (Figs. 4B, S6Cand Supplementary Information). Compared to *C-GAP* RNAi embryos, the predicted wild-type apical cell area profile is far more homogeneous (cf. Figs. 3D, 4C), and averaging over 50 realizations of the myosin fluctuations showed that stochastic pulsing considerably lowers variations in apical areas (Fig. 4E). With pulsing, normal furrowing is recovered, the predicted furrow shape matches experiment (Fig 4C) and the furrow shape failure rate is zero (Fig. S7A).

How does myosin pulsing rescue tissue contraction and furrowing? Since the myosin level determines the contraction rate, the apical cell area profile is set by the time averaged myosin profile. Now without pulsing relative myosin fluctuations are unchanging, so time-averaging has no effect and tissue contraction is severely disrupted. By contrast, time-averaging the pulsatory fluctuating myosin signal progressively lowers its relative fluctuations, reducing their disruptive impact on tissue contraction. Equivalently stated, by cycling through the range of accessible values, pulsing effectively eliminates high frequencies in the myosin signal, reducing the toal fluctuation power (Fig. 4F). Thus, pulsing is a low pass filter mechanism that rescues tissue contraction and folding.

### Twist depletion disrupts furrowing due to the absence of an envelope and large myosin fluctuations

Our results suggest: (i) furrowing is driven by curvature in the myosin envelope, and (ii) tissue contraction is highly sensitive to cell-to-cell myosin fluctuations, so even when a cell recruits active myosin it will fail to contract if its myosin level is persistently below that of its neighbors (Fig. 3A). We now show that these two principles explain the experimentally observed failure of VFF in embryos depleted of Twist, a transcription factor whose downstream genes activate the RhoA pathway leading to myosin ramp-up^3^.

In *twist* RNAi embryos, ROCK and myosin fail to ramp up, and in a region within ∼ 13 cells of the ventral midline cells instead exhibit asynchronous pulses of myosin on a zero baseline (Fig. 5A)^21^, presumably induced by the transcription factor Snail that is uniformly distributed over the presumptive mesoderm^50^. Apical cell areas are highly inhomogeneous, some cells contracting and others expanding with no overall tissue contraction^10,21^, and the ventral surface fails to furrow^5^ (Fig. 5C, right).

**Figure 5.**
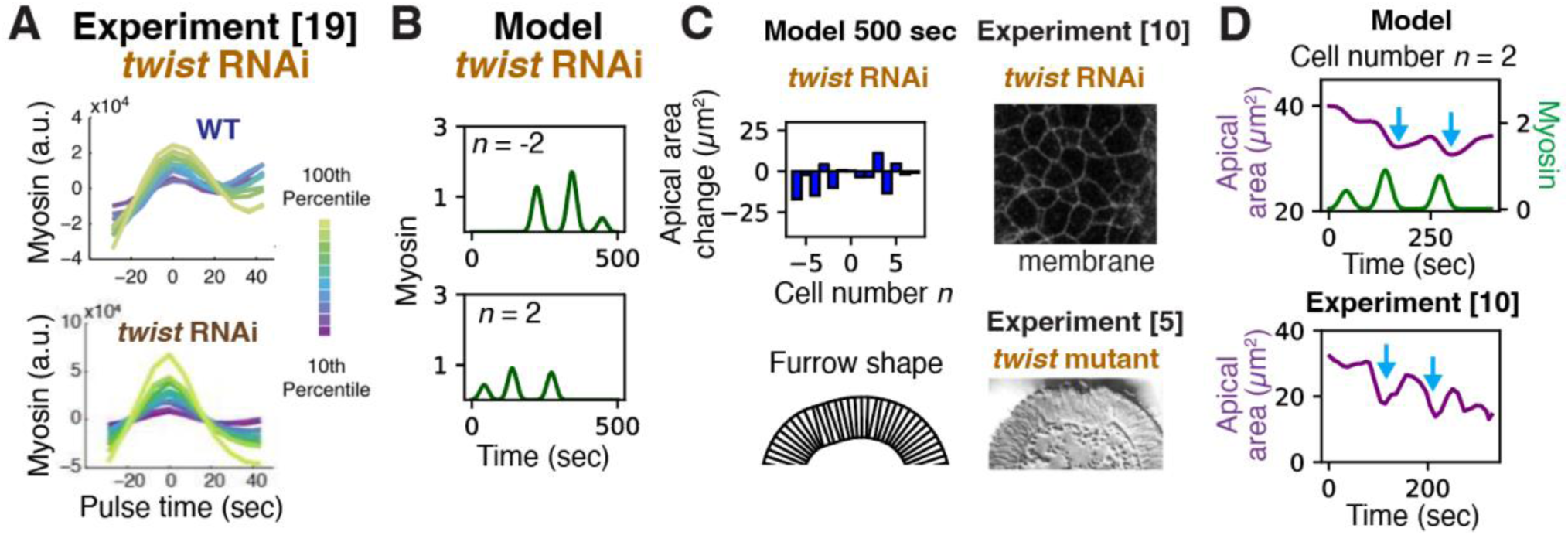
Twist depletion abolishes tissue contraction and furrowing. (A) In Twist-depleted embryos, myosin pulses return to baseline without net ramp-up. Experimentally measured mean myosin vs. time during individual pulses in different amplitude intervals (n > 80 for each wild-type curve, n ≥ 110 for each *twist* RNAi curve). Adapted with permission from ref.^21^. (B) Typical myosin profile used as input to model *twist* RNAi embryos, in which myosin pulses stochastically without ramp-up. Pulse statistics are taken as the experimental statistics of (A). (C) Predicted apical area profile and tissue shape at 500 sec (left). Some cells contract while others expand, but no net contraction or furrowing is accomplished. Fluorescence micrograph of the presumptive mesoderm in a *twist* RNAi embryo (top right) and transverse view elctron micrograph in a *twist* mutant (bottom right), adapted with permission from refs.^10^ and ^5^, respectively. Consistent with model predictions, cells variably contract or expand without net contraction, and furrowing fails. (D) Apical myosin and model-predicted apical area vs. time for a typical cell (top). Experimental apical area vs. time for a *twist* RNAi embryo, adapted from ref.^10^ with permission. Blue arrows, “unratcheted” contraction pulses.

To model *twist* RNAi embryos, we mimic the myosin signals in individual cells by a series of 35 s long^21^ pulses on a zero baseline for cells -6 to 7, with random intervals from 30s to 330 s between pulses and random pulse amplitude from zero to twice the wild-type myosin amplitude (see Fig. 2) (Fig. 5B). In agreement with experiment^5,10,21^, the model predicts that some cells expand while others contract, with zero mean contraction and hence no furrowing (Figs. 5C, S7B). In some cases, contracting cells subsequently expand (blue arrows) as seen experimentally^21^ (Fig. 5D).

These results suggest furrowing failure in *twist* RNAi embryos originates in the absence of an envelope. Without Twist-dependent ramp-up, the myosin envelope whose curvature normally drives furrowing is absent, and myosin pulses occur against a zero baseline so that cell-to-cell relative fluctuations are ∼ 100%, provoking such large apical area fluctuations that many cells fail to contract.

## Discussion

The programs of development are extraordinarily robust to fluctuations, leading to the same adult organism with relatively little phenotypic variation despite stochasticity in the genotype^51,52^, in the environment^53,54^ and at the cellular and multicellular levels^22,23^. At cellular and tissue levels, development relies on signalling cascades whose ability to transform information from one form to another is limited by intrinsic molecular stochasticities^24^. In the classic case, concentration profiles of morphogen proteins encode gene expression profiles^55,56^. For example, in the scyncitial *Drosophila* blastoderm the concentration profile of the Bicoid (Bcd) morphogen helps specify the spatial expression profile of the gene that encodes hunchback (Hb), which together with other gap genes will determine anterior-posterior segmentation^57-59^. In ref.^60^ it was argued that the spatial resolution with which the Hb profile is specified lies just within the limits imposed by fluctuations in local Bcd concentrations, while other studies suggest stochasticities are controlled by cross-regulation among gap genes^61-63^. In other signalling contexts noise may be controlled by negative feedback loops^24,64^.

In another class of signalling cascade, spatial protein distributions are transformed into mechanical output. The concentration profile of a motor protein, say, can encode tissue morphology, and its ability to do so may be noise-limited. During *Drosophila* VFF, the profile of apical myosin levels among cells encodes ventral epithelial tissue shape. We concluded that apical cell surface contraction rates and tissue shape are specified by curvature in the myosin profile and the anisotropic curvature of the envelope of the profile would encode a perfect furrow with anterior-posterior orientation (Fig. 2C). However, there are substantial fluctuations about this smooth envelope. Generally, fluctuations in protein copy numbers are unavoidable due to multiple upstream processes, including transcription and translation, gene regulatory networks, protein-activating signalling cascades, and diverse cellular events such as trafficking and exocytosis^22-24^. The stochasticity of these microscopic processes originates in the small copy numbers of genes, RNA transcripts, proteins and other molecules involved. In the case of *Drosophila* VFF, activated apical myosin levels have ∼ 45% cell-to-cell fluctuations about the envelope (Fig. S6)^12-14^, reflecting a cascade of stochastic upstream processes starting with the dorsal-ventral patterning morphogen Spätzle, whose profile is set by asymmetric maternal modification of the vitelline membrane. The Spätzle profile determines expression patterns of the transcription factors Twist and Snail over ventral cells^3^, which in turn encode the expression profiles^47^ of the signaling molecules Fog, the GPCR Mist and T48. Secreted Fog binds the Mist receptor (itself dynamically regulated by exocytosis/endocytosis^48^) and both Fog-bound Mist and T48 recruit RhoGEF2 which activates the RhoA signaling pathway^65,66^. This leads to the activation of formin and ROCK, which respectively nucleate actin filaments and activate myosin^9^.

Are these substantial fluctuations sufficient to wash out the mechanical information encoded in the myosin profile? Surprisingly, we found the answer is yes, but only if the fluctuations persist in time. We found a ∼ 25-55% defecit myosin fluctuation relative to the envelope is sufficient to prevent a cell from contracting, Eq. (3) (Fig. 4A). Since actual fluctuations are of this order or greater^12,14^ (Figs. 3, S6A, B), we expect severe disruption of furrowing. This is indeed seen in *C-GAP* RNAi embryos, when myosin ramps up at stochastically different rates in different cells, but the rate in a given cell is approximately fixed so fluctuations relative to the envelope are locked in. In these embryos, and in *sqh-EE* mutant embryos for similar reasons, some cells fail to constrict and furrowing is impaired^38,39^ (Fig. 3). Similarly, in a study using optogenetic RhoGEF2 to activate apical myosin in the *Drosophila* germband, myosin levels fluctuated among cells, and cells variably contracted or expanded, with no collective contraction^67^. This may result in a situation similar to that in *C-GAP* RNAi embryos during VFF, with high RhoGEF2 levels but low RhoGAP; indeed, compared to wild-type VFF, myosin fluctuations were temporally persistent^67^. Relative myosin fluctuations are also extremely high in *twist* RNAi embryos since the envelope is absent, so cells fail to contract and furrowing fails^5,10^ (Fig. 5).

In wild-type embryos, these disastrous effects are rescued by stochastic pulsing of myosin (Fig. 4A, B) whose origin is thought to be cyclic activation and deactivation of RhoA by RhoGEF2 and C-GAP in the RhoA pathway upstream of myosin^49^. In contrast to *C-GAP* RNAi embryos, myosin is cycled every ∼ 1 min through the full fluctuation range about the monotionic ramp-up, so the signal is time-averaged. High frequencies in the fluctuation power spectrum are thus eliminated (Fig. 4F), a low-pass filter mechanism reducing net fluctuations sufficiently that ventral cells contract homogeneously and furrowing is rescued (Figs. 4, 6). A necessary feature for this mechanism to work is that cells pulse independently, so a cell’s myosin is sometimes above and sometimes below that of its neighbors. Indeed, asynchronous stochastic cycling is presumably far more easily accomplished than synchronized behavior. We suggest this is a broadly employed method to control fluctuations, since myosin pulsing is a common feature of many contractility-driven morphogenetic processes^26-31,68^. Other examples in *Drosophila* are the junctional myosin pulses during germband extension^29,30^ and apical myosin pulsing in amnioserosa cells during dorsal closure^33,34^, where the much slower pulsing (∼ 3-4 min) is consistent with the longer ∼ 40 min duration of the fast phase. Actomyosin pulsing is also a feature of *C. elegans* gastrulation^35^ and *Xenopus* neurulation^36^.

**Figure 6.**
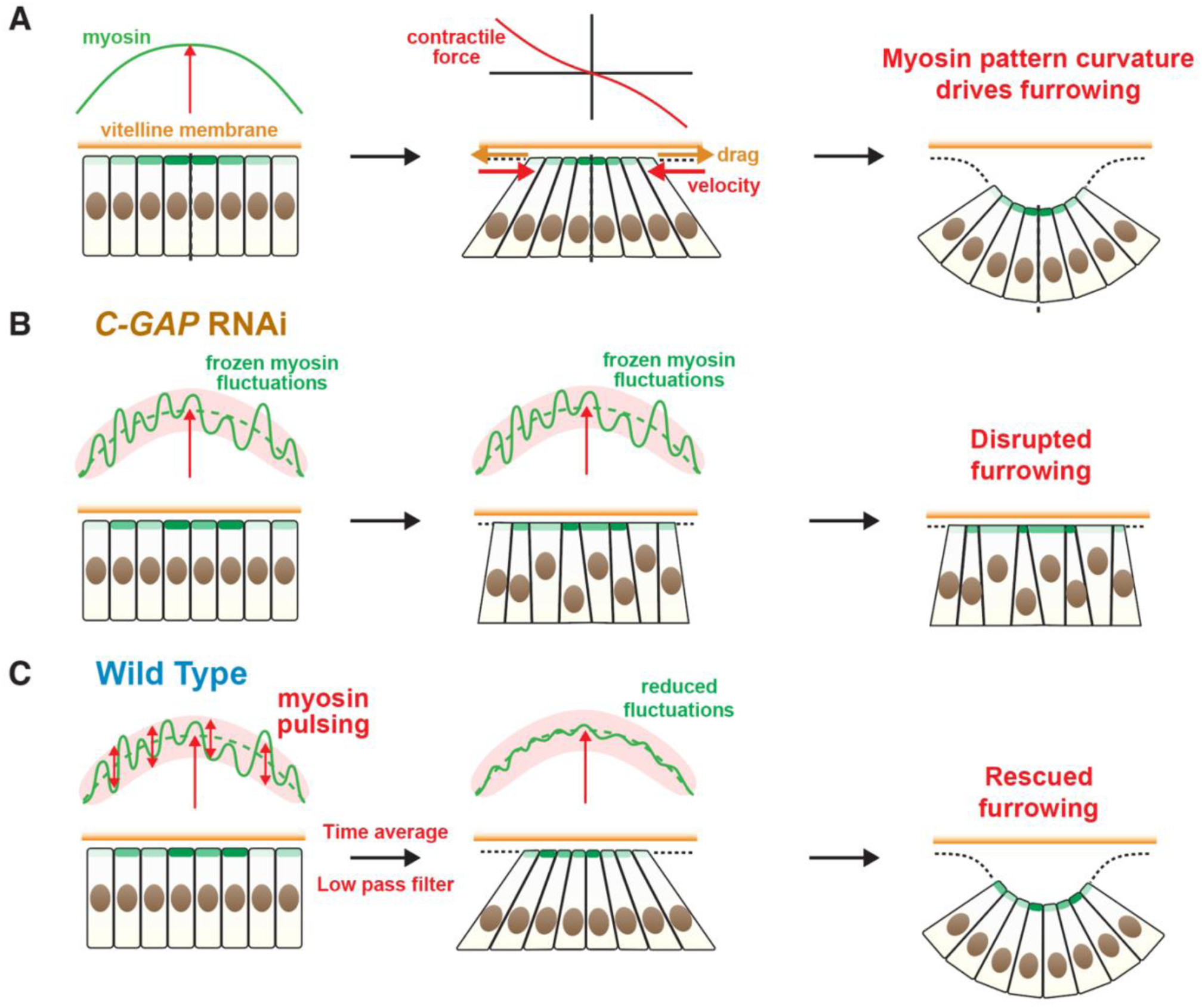
Model of *Drosophila* ventral furrow formation. (A) Curvature in the myosin profile drives furrowing. The furrow curvature is created by constriction of the apical surfaces of rows of cells straddling the ventral midline, requiring that cells be moved relative to the vitelline membrane by actomyosin contractile force. This motion is opposed primarily by frictional drag from the vitelline membrane; the elastic resistance is weak, since the stiff nuclei move toward the basal cell surface. Since cells further from the midline move faster the drag force force increases with distance from the midline, so the same must be true of the contractile force that generates this motion. Since the contractile force is proportional to the gradient of the myosin profile, the myosin profile must have curvature. (B) In *C-GAP* RNAi embryos, relative myosin fluctuations are frozen in time as the profile ramps up (red vertical arrow). Apical cell surface contraction is highly sensitivity to these cell-to-cell fluctuations, so tissue contraction is inhomogeneous and furrowing is impaired. (C) In wild-type embryos, myosin levels in cells pulse asynchronsously and stochastically in time about the smooth envelope. The time-averaged signal in each cell has reduced amplitude, a low pass filter mechanism that rescues furrowing.

A central conclusion of this study is that ventral reshaping is encoded in curvature of the myosin patterning. Since curvature is in general direction-dependent, diverse tissue shapes can in principle be sculpted. During VFF, the myosin envelope has curvature in the ventral-lateral direction but is flat over more than 30 cells along the anterior-posterior axis, so a furrow oriented in the anterior-posterior direction is sculpted^14^ (Figs. 2C and S3). This explains why cells constrict apically in the ventral-lateral direction but not in the anterior-posterior direction^7,40^, a behavior inconsistent with elastic models. The curvature requirement also explains furrowing failure in Spn27A depleted embryos, whose top-hat profile is flat in the central activated zone (Fig. 2A).

The significance of the myosin profile curvature originates in the dominant role of frictional drag forces in resisting myosin contractility, because apical constriction of a row of cells spanning the activated myosin zone requires their displacement toward the midline (Fig. 1A). Cells further from the midline require greater displacement, must translate with higher velocity, and hence encounter greater frictional drag (Fig. 1A, 6A). Thus, the actomyosin force propelling the cells must also increase with distance from the midline. Since the actomyosin force is proportional to the myosin gradient, the myosin profile must have curvature.

Previous modeling analyses of *Drosophila* VFF primarily assumed elastic tissue response^11,16^ and failed to explain basic observations such as the localized ∼ 2-3 cell-wide expansion zones just outside the central contraction zone, or non-contraction of ventral cells in the anterior-posterior direction. Indeed, we found that adding substantial elastic forces to our model describing the myosin-mediated collective cell contraction led to growing expansion zones inconsistent with experiment (Fig. S4). We argued that elastic stresses are far too small to resist the forces exerted by myosin. To induce epithelial tissue curvature for furrowing, *Drosophila* constricts apical cell surfaces using actomyosin networks. Similar actomyosin networks were recently measured by magnetic tweezers to exert forces *T*_myo_ ∼ 1-1.5 nN during *Drosophila* dorsal closure^69^. Elastic resistance cannot compete with such forces: the elastic constant for apical cell deformation was measured to be *k* = 7 pN μm^−1^ using magnetic beads during late *Drosophila* cellularization^19^, so even for complete apical constriction (cell width decrease Δ*L* ∼ 7 μm), the elastic resistance *k* Δ*L* ∼50 pN is over 20-fold smaller. Indeed, apical constriction is a very particular and soft cell deformation mode (Fig. 1A) in which the basal cell surface expands and the very stiff nucleus shifts toward the basal surface and avoids compression^70^. Generally, on account of its high stiffness, the nucleus can be a determinant of cell and tissue shape evolution in response to forces^71^.

Thus, we conclude that apical cell constriction is resisted primarily by viscous forces during VFF. This conclusion is qualitatively consistent with a study showing that tissue deformation during *Drosophila* VFF can be described as a viscous fluid flow system^72^. Another key study of *Drosophila* VFF and germband extension accounted for viscous forces to predict tissue velocity fields^73^, but tissue reshaping was not predicted. Here, we fitted model-predicted apical area profiles and contraction rates to experiment for best fit values of the the physical constants *λ* and *μ* governing external and internal viscous forces, respectively (Table 1). These values are much higher than those measured during cellularization^19^, as expected since after cellularization ventral cells are pushed against the vitelline membrane^45^ and apical networks assemble^10^. External drag forces resisting apical cell surface motion are likely dominated by friction with the inner eggshell layer, the vitelline membrane^19^, while the cell viscosity *μ* = 36 pN s nm^−1^ measures dissipative effects associated with apical area change including apical network remodeling requiring dissipative motion of protein complexes anchoring the network to the apical membrane. The associated viscous forces are unknown, but related properties were measured in the cytokinetic actomyosin contractile ring that divides fission yeast cells^74-77^, anchored to the plasma membrane by protein complexes called nodes. Each node has a drag coefficient ∼ 0.5 pN s nm^−1 78^, so during *Drosophila* VFF the apical cell network offers the resistance of ∼ 70 fission yeast nodes.

## Materials and Methods

### Mathematical model of VFF

To drive ventral issue contraction in the *Drosophila* blastoderm, apical actomyosin networks generate contractile stresses on the apical cell surfaces of variable magnitude from cell to cell, whose magnitudes are assumed proportional to the amount of activated myosin II in that cell (Fig. 1A, B, C). We follow the standard notation: the ventral midline extends along the length of the embryo, the anterior-posterior axis, while the ventral-lateral axis orients perpendicular to that direction (Fig. 1C). On the embryo ventral surface, the myosin pattern follows a ∼ 10-15 cell wide bell-shaped profile along the ventral-lateral axis, but is approximately flat in the anterior-posterior direction, extending ∼ 30 cells in that direction with constant amplitude for a given ventral-lateral location. Indeed, cell properties are statistically independent of anterior-posterior axis location over ∼ 30 cells, including apical areas, contraction rates, velocities and apical myosin levels^6,7,14^. Thus, for simplicity we consider variations in the ventral-lateral direction only, with *n* labelling the *n*^th^ cell from the ventral midline (Fig. 1C). At the location of maximum width of the ellipsoidal *Drosophila* blastoderm, ∼ 80 connected cells extend along the ventral-lateral axis^10^.

The envelope of the myosin II pattern is bell-shaped and ramps up over ∼ 7 min in wild type embryos^12-14^. Equally important, there are large fluctuations about this smooth envelope, both spatially (from cell to cell) and temporally (pulsatile time dependence within a given cell)^12,14^. When the contiguous cells are subject to these actomyosin contractile stresses, cell translation is opposed by external drag forces likely due to the vitelline membrane^19^ (Fig. 1D) and apical cell contraction is opposed by internal viscous forces, proposed to originate from microtubules^19^ and the cytoplasm^79^ during cellularization before apical network formation. During VFF, internal dissipation may be dominated by anchoring and other effects associated remodeling the apical networks. Finally, elastic forces also resist apical constriction^19,79^.

Let *L*(*n*) and w(*n*) denote the width and contraction rate of cell *n* in the vental-lateral direction. The total cell tension is *T*_tot_(*n*) = *T*_myo_(*n*) − *μw*(*n*) − *k*Δ*L*(*n*) where *T*_myo_(*n*) is the actomyosin contractile tension, *μ* the internal viscosity, *k* the elastic force constant and Δ*L* the width relative to the resting width (Fig. 1D). The force balance on the cell is *T*_tot_(*n* + 1) − 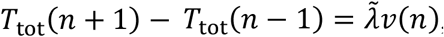, where 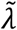 is the external drag coefficient per cell and *ν*(*n*) the cell velocity. Thus,

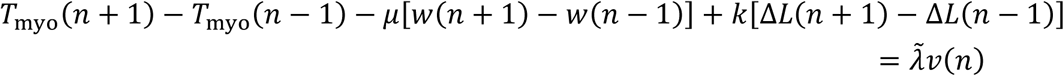

Defining 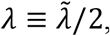, in the continuous limit this gives Eq. (1) of the Model section.

We now argue the elastic term is small and can be discarded. A recent magnetic tweezers study measured single cell apical myosin contractile stresses *T*_myo_ ∼ 1-1.5 nN during *Drosophila* dorsal closure^20^. For the elastic constant we use *k* = 7 pN μm^−1^ measured in the experiments of ref.^19^ during late cellularization, when magnetic beads were manipulated to provoke apical deformation similar to that during VFF. Note this is a very particular deformation mode, with contraction near the apical surface only and nuclei being pushed downwards^70^. Taking a typical cell width 7 μm^10^, it follows that even for 100% strain the internal elastic stress *k*Δ*L* ∼ 50 pN is far smaller than the above estimated myosin contractile stress. The elastic term can thus to a good approximation be neglected in Eq. (1). Taking the derivative with respect to *n* and noting *w* ≡ −∂*ν*/∂*n*, we obtain Eq. (2) of the Model section.

Throughout this study, we use experimental data to obtain the myosin pattern *T*_myo_(*n, t*). Using this in Eq. (2), we solve for the contraction rate *w*(*n, t*), whose time integral equals the net cell contraction profile and hence the apical width profile *L*(*n, t*) which is proportional to the apical area profile assuming fixed cell lengths in the anterior-posterior direction (Fig. 1A, B). In cases where a cell contracts to zero width, the cell is removed from the dynamics.

### Determination of model parameters by fitting experiment

Solutions to Eq. (2) involve just two reduced parameters: the damping length *ξ* = (*μ*/*λ*)^1/2^, and the characteristic contraction rate 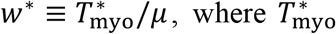, where 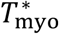 is the maximum amplitude of the myosin profile. For wild type embryos we take 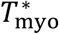 to be the maximum of the myosin tension profile at 300 s (the value at the ventral midline). (1) Using the myosin levels measured in ref.^12^ for cells 1-10, we compared the model-predicted apical cell area profile to that measured in the same study. This yielded a best fit value *ξ* = 2.5 ± 1.2 (Fig. S1). (2) Using the myosin levels and contraction rates of cells 1-7 at ∼ 300 s measured in ref.^14^, a best fit of the predicted contraction rate profile yielded *w*^*^ = 42 ± 6 nm s^−1^ (Fig. S1). (3) Using 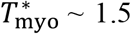 nN from ref.^20^ and the above values of *ξ* and *w*^*^, we obtain estimated values for the external drag coefficient *λ*∼6 pN s nm^−1^, and for the intracellular viscosity *μ*∼36 pN s nm^−1^ (Table 1). For details, see Supplementary Information.

### Furrow shape calculation

Contraction of apical cell surfaces generates curvature and furrows the ventral surface (Fig. 1A). From the predicted apical area profile, we calculated the multicellular tissue shape assuming the apical and basal cell areas are in the ratio of their respective radii of curvature, and that the cell height and volume are conserved and equal to 39 μm^80^ and ∼ 1500 μm^3 70^, respectively. This uniquely determines the shape of each cell. The cells were then joined to obtain the cross-sectional tissue shape. For details, see Supplementary Information and Fig. S2.

### Two dimensional model of ventral furrowing

To test our analysis that assumed variations in the ventral-lateral direction only, we developed a 2D analysis explicitly accounting for dependencies in the anterior-posterior direction. The contraction rate *w* ≡ −*dν*/*dn* is generalized to *w* ≡ −**∇**_**n**_ · **v**, where **n** is the cell number vector. The myosin tension *T*_myo_ in Eq. (2) is generalized to an isotropic myosin stress **σ**_myo_ = *σ*_myo_**I**. The internal viscous force *μw* generalizes to an isotropic stress *μw***I** = −*μ*(**∇**_**n**_ · **v**)**I**. The balance of myosin, internal viscous and external drag forces reads

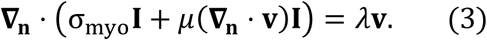

Using the 2D myosin envelope taken from experiment^15^, this system was solved numerically using a Fourier spectral method to obtain the velocity and contraction rate fields, **v**(**n**) and *w*(**n**). The predicted contraction rate profile agrees with the 1D analysis (see Fig. S3).

### Tissue contraction solutions for hypothetical myosin profiles with and without curvature

In addition to actual experimental myosin profiles, we solved Eq. (2) for two hypothetical profiles lacking curvature (top-hat and triangular), and one profile possessing curvature (parabolic) (Fig. 1E). The exact solutions for the contraction rate profiles are (see Supplementary Information)

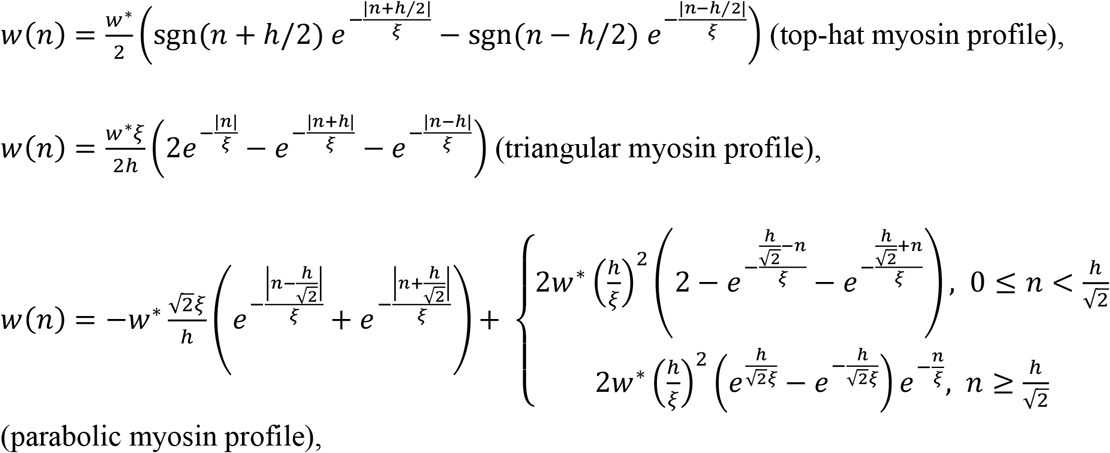

where sgn(*x*) ≡ *x*/|*x*|. Here *h* is the width of each profile (number of cells) and 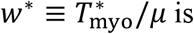 is a characteristic contraction rate, where 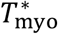 is the maximum amplitude of the myosin stress profile. Since the myosin patterning was taken as time-independent, the same is true of the above contraction rate profiles *w*(*n*). Thus, relative to the resting value with no myosin, the profiles of changes in cell widths and changes in cell areas after time *t* are, respectively, Δ*L*(*n*) = −*w*(*n*) *t* and Δ*A*(*n*) = −*b w*(*n*) *t* where *b* = 5.7 μm is the effective initial cell length in the anterior-posterior direction^79^. Thus the cell area profiles are proportional to the contraction rate profile expressions above. The three hypothetical myosin patterns and the apical area profiles they generate are shown in Fig. 1E for times of several hundred sec, with the corresponding furrow shapes calculated using the method outlined above.

These results show that activated myosin is in itself insufficient to collectively contract the ventral cells: even in the high amplitude regions near the ventral midline, if the myosin pattern has constant amplitude (top-hat profile) or constant gradient (triangular profile) little cell contraction occurs and a severely misshapen furrow results (Fig. 1E). For these profiles, constriction occurs only near (within distance *ξ* of) singular points with infinitely negative curvature (top hat edges or triangle vertex). By contrast, when the myosin profile has curvature (parabolic profile) extended cell contraction occurs, producing a nicely rounded furrow.

The origin of this requirement is that apical constriction of contiguous cells requires cell translation, with cells more distant from the ventral midline having higher velocity (see Fig. 1A). On the large scales of the myosin envelope, this motion is primarily resisted by frictional drag forces from the vitelline membrane and other external entities. Thus, to move the cells in this manner requires the actomyosin contractile force acting on a cell to be greater for cells more distant from the midline. Since the contractile force varies as the gradient of the myosin level, it follows that the myosin profile must have curvature. This is explicit from Eq. (2) which, for scales much larger than the damping length *ξ* ∼ 2-3 cells, simplifies to *λw* ≈ − ∂^2^*T*_myo_/∂*n*^2^, i.e., the contraction rate is proportional to the myosin envelope curvature. This point is discussed further in Discussion.

### Modeling spatiotemporal myosin fluctuations in wild-type and *C-GAP* RNAi embryos

To explore the effects of temporal myosin fluctuations in wild-type and *C-GAP* RNAi embryos, we mimicked the experimental temporal myosin profiles. We solved Eq. (2) using as input time- and cell-dependent myosin profiles of the following form:

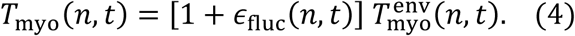

Here 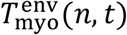 is the envelope for wild-type embryos, whose amplitude increases linearly in time. The relative fluctuation factors *ϵ*_fluc_(*n, t*) for different cells are statistically independent, and their time dependencies are chosen to mimick the experimental myosin signal in wild-type (pulsatile fluctuations) and *C-GAP* RNAi (persistent fluctuations) embryos (Figs. 3B, 4C) with different autocorrelation timescales. See Supplementary Information and Fig. S6C for details. Results of these calculations are shown in Figs. 3D, E, and Figs. 4C-E.

## Acknowledgements

This work was supported by National Institute of General Medical Sciences of the National Institutes of Health under award number R01GM086731 to B.O’S. The content is solely the responsibility of the authors and does not necessarily represent the official views of the National Institutes of Health.

## Author contributions

B.O’S. conceived the study. B.O’S. and H.Z. designed the model and performed the mathematical modeling. H. Z. numerically solved the equations. B.O’S. and H.Z. analyzed the data. B.O’S. and H.Z. wrote the manuscript.

## Competing financial interests

The authors declare no competing financial interests.

## Data and Code Availability

Simulation results supporting the findings of this paper, and codes to perform the simulations, to analyze the data, and to generate the technical figures are available in the GitHub repository [https://github.com/OShaughnessyGroup-Columbia-University/drosophila_ventral_furrow_formation.git].

## Notes

### Competing Interest Statement

The authors have declared no competing interest.

### Summary of Updates

Format updated.

