## Supplementary Information for "Actomyosin pulsing rescues embryonic tissue folding from disruption by myosin fluctuations"

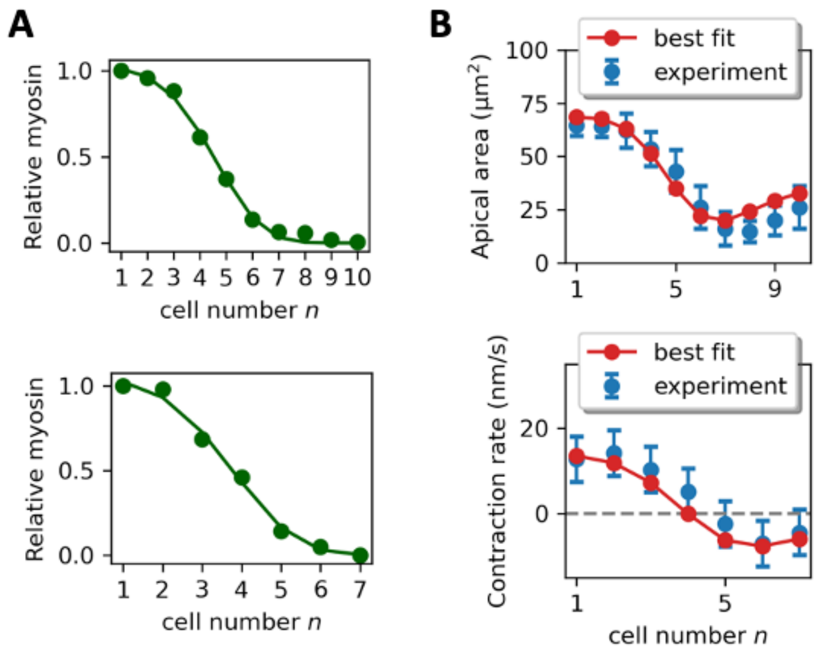


**Figure S1. Determination of best fit values of model parameters $\xi$ and $w^*$ from experimentally measured cell apical area and contraction rate profiles.** (A) Smooth myosin envelopes fitted from experimental myosin levels in ref.[1] (top) and ref.[2] (bottom), used as input for fitting model parameters $\xi$ and $w^*$. (B) Top: experimental apical areas of cells 1-10, taken from ref.[1]. Best fit model prediction is shown (red) using the damping length $\xi$ as fitting parameter (best fit value is $\xi = 2.5 \pm 1.2$). Bottom: experimental contraction rates of cells 1-7, ~ 300 sec after gastrulation onset, taken from ref.[2]. Best fit model prediction is shown (red) with the value $\xi = 2.5 \pm 1.2$ and using the characteristic contraction rate $w^*$as fitting parameter (best fit value is $w^* = 42 \pm 6\ \mathrm{nm/s}$).

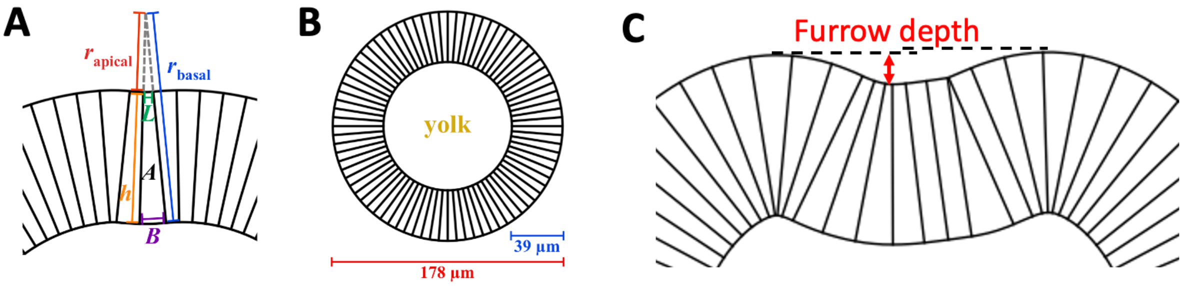


**Figure S2. Determination of tissue shape for a given apical area profile, and the definition of furrow depth.** (A) For each cell we assume the ratio of the apical radius of curvature $r_{\mathrm{apical}}$ and the basal radius of curvature $r_{\mathrm{basal}}$ equals the ratio of the cell's apical width $L$ to its basal width $B$. In addition, we assume the apico-basal length $h$ is fixed at 39 µm for all cells, according to the measured apico-basal length ~ 30-40 µm during ventral furrow formation[3]. We also assume the volume of all cells is fixed at 1500 $\mu m^3$ as measured experimentally[4]. This translates to a cross-sectional area of $A = 214\ \mu\mathrm{m}^2$ for each cell. Therefore, for each cell $n$ at time $t$, the apical width $L(n)$ is obtained, and together with the apico-basal length $h$ of 39 µm and the cross-sectional area $A$ of 214 $\mu m^2$, the basal width $B(n)$ and the apical and basal radii of curvature $r_{\mathrm{apical}}(n)$ and $r_{\mathrm{basal}}(n)$ are solved for. We then join the cells together by connecting their apical and basal surfaces to obtain the complete tissue shape. (B) The initial blastoderm before gastrulation assumed by the present model. Experimentally[3], the blastoderm diameter is ~ 180 µm. (C) The furrow depth is defined as the height difference between the trough of the furrow and the lower crest (indicated by the red double arrow).

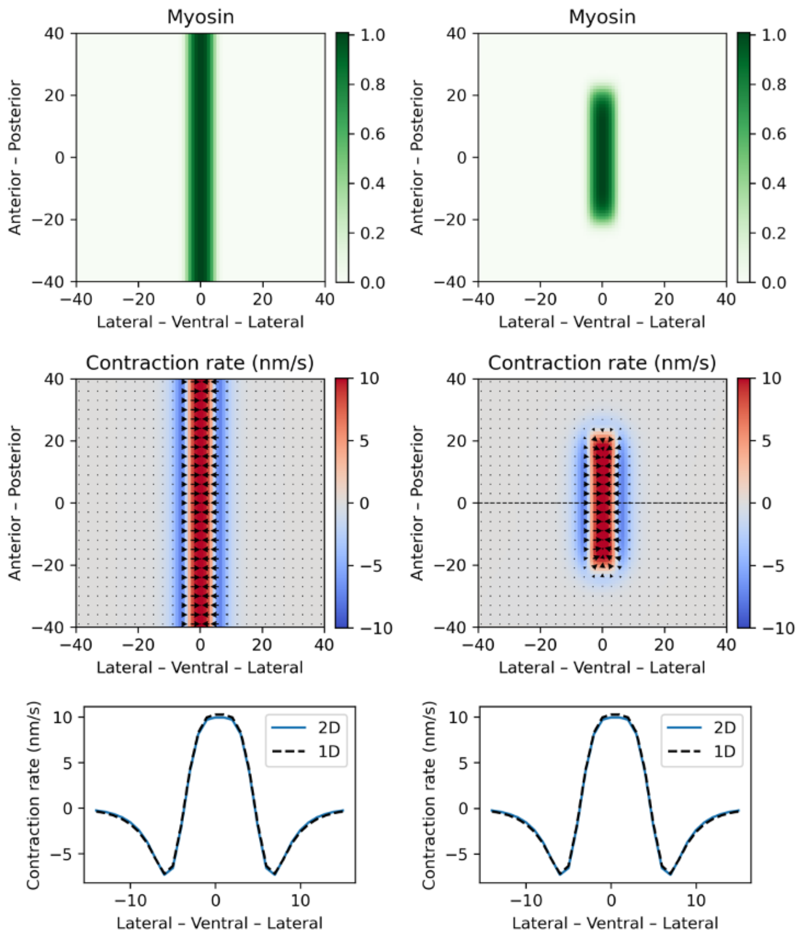


**Figure S3. Predictions of 2D continuum model.** The embryonic tissue is represented as a sheet of 80 x 80 cells. Contraction rates are calculated by solving Eq. 3 in Materials and Methods. Left column: A myosin envelope with amplitude independent of anterior-posterior position is used as input to the model (top). The predicted contraction rate profile (colors) and velocity profile (arrows) is independent of anterior-posterior location (middle), and the contraction rates agree with the profile predicted by the 1D model presented in Fig. 2A (bottom). Right column: results for a more realistic experimental myosin envelope extending ~ 30 cells in the anterior-posterior direction axis (top). The predicted contraction rate profile (colors) and velocity profile (arrows) are independent of anterior-posterior location from approximately cell −10 to cell +10 (middle). Predicted contraction rates along the dashed line agree with the 1D model in Fig. 2A (bottom).

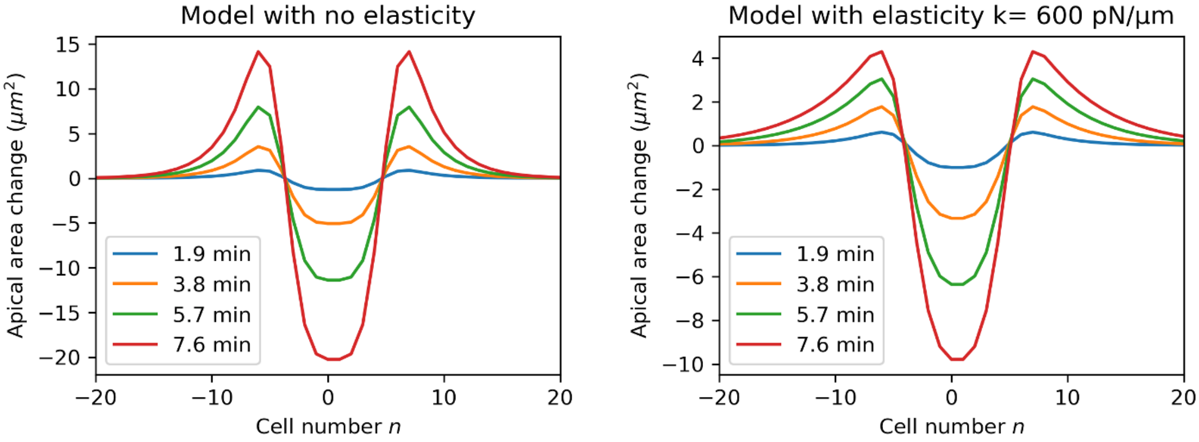


**Figure S4. A model assuming substantial elastic forces fails to reproduce the experimentally measured cell apical area profiles during *Drosophila* ventral furrow formation.** Cell apical area profiles predicted by the model presented in this work (Eq. (2)), incorporating intracellular viscous forces and extracellular drag forces parametrized by internal viscosity $\mu$ and external drag coefficient $\lambda$, respectively (left). This model reproduces the experimental observation[5] that two fixed bands of 4 cells each suffer expansion during VFF, located lateral to the region of maximal myosin activation (cells ~~5~~ 6 to 8 and -~~5~~ 6 to -8). In contrast, when elastic forces are added into this model with elastic constant $k = 600$ pN/μm, the number of cells in the predicted expansion zones continues to increase with time throughout VFF (right).

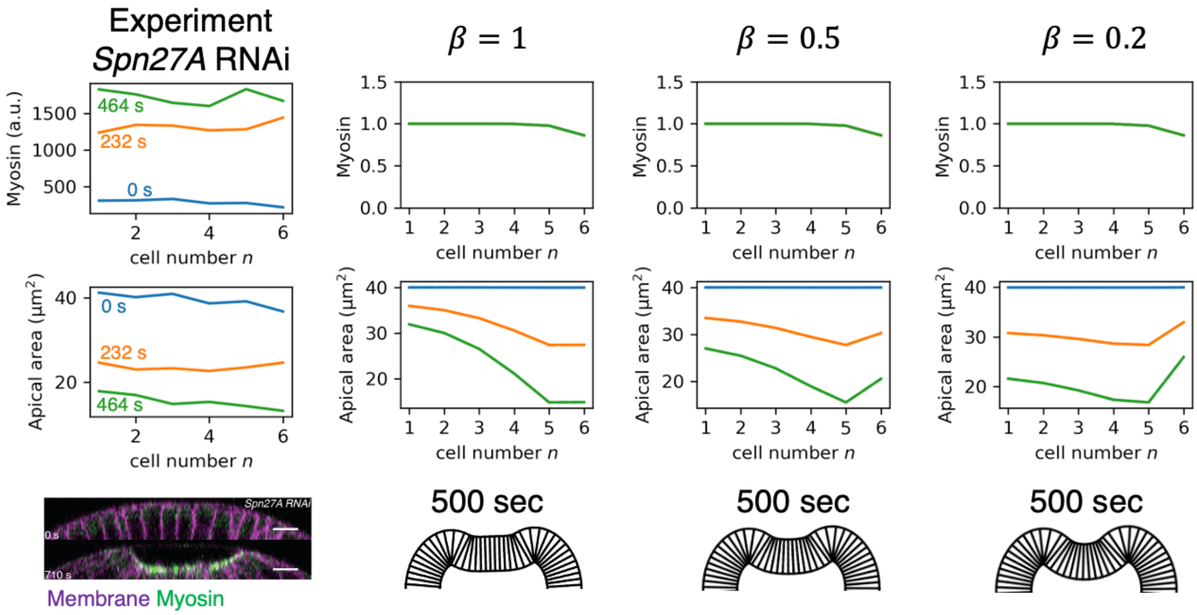


**Figure S5. Predicted cell apical area profiles and furrow shapes in *Spn27A* RNAi embryos for three values of the external drag reduction factor $\boldsymbol{\beta}$.** Left column: experimental myosin profile, apical area profile and furrow shape in *Spn27A* RNAi embryos, adapted with permission from ref.[2]. Right: each column shows the predicted for apical area profile and furrow shape for *Spn27A* RNAi embryos using the plotted myosin profile as input (same for all columns). For each column a different value of $\beta$ was used, where the model used an external drag coefficient equal to $\beta\lambda$ for cells -6 to 6 (wild-type case is $\beta = 1$).

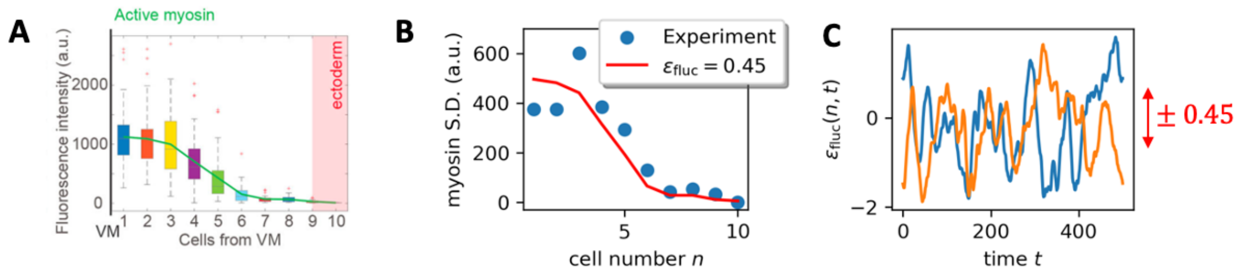


**Figure S6. Extracting magnitudes of spatiotemporal fluctuations in myosin levels from experimental measurements in wild-type cells (related to Fig. 6).** (A) Box-and-whisker plot of experimentally measured myosin levels for cells 1-10, adapted with permission from ref.[1]. (B) Standard deviations of myosin level versus cell number obtained from (A) (blue points). Standard deviation profile assuming fluctuations relative to the mean value have a fixed value $\epsilon_{\text{fluc}}$ ($\epsilon_{\text{fluc}} = 0.45$ is the best fit using non-linear fitting) for all cells (red curve). (C) Two realizations of the stochastic factor $\epsilon_{\text{fluc}}(n, t)$ that generates the fluctuating wild-type myosin signal of Fig. 4. The variation amplitude is ± 0.45, as extracted in (B).

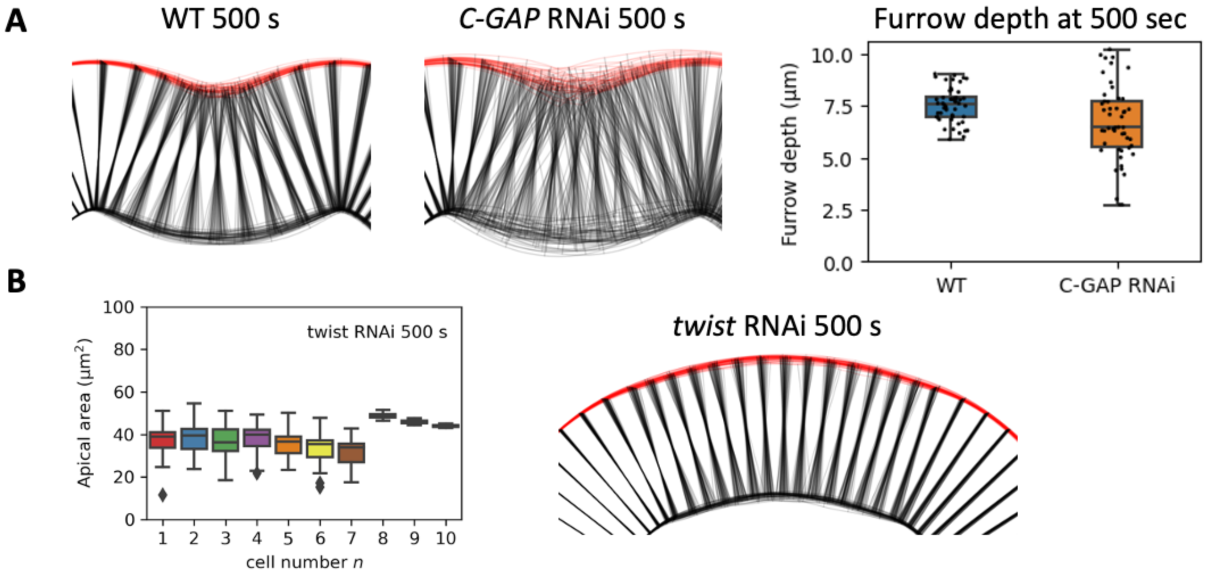


**Figure S7. Furrow shapes and apical area profiles predicted by model, showing mean values and fluctuations.** (A) Predicted furrow shapes for wild-type and *C-GAP* RNAi embryos. For each case, predictions from 50 independent solutions are superposed. Apical surfaces shown in red; lateral and basal surfaces shown black. Predicted fluctuations in furrow shape are much higher for *C-GAP* RNAi embryos than for wild-type. Furrow failure is defined as the furrow depth smaller than 60% of the furrow depth without myosin fluctuations. (B) (Left) Box-and-whisker plots of model-predicted apical area profiles at 500 sec for *twist* RNAi embryos (50 simulations). (Right) Predicted tissue shapes for *twist* RNAi embryos (50 predictions superposed) show that ventral furrows do not form in *twist* RNAi embryos, consistent with experiment.

### Using the model to predict the cell apical area profile

From the model equation Eq. (2), we predict the contraction rate $w(n,t)$ for each cell based on the myosin profile $T_{\text{myo}}(n,t)$. This contraction rate is the rate of change of the apical width of the cell $L(n,t)$, giving the apical length profile $L(n,t)$,

$$L(n,t) = L_0 - \int_0^t w(n,t)dt\,.$$

Initially the epithelial sheet of cells is assumed uniform, with cell apical width $L_0 = 7\ \mu\text{m}$, consistent with the cell diameter before gastrulation[6]. Based on the apical width profile, we predict tissue shape using the algorithm described in Fig. S2. Using the fact that contraction in the anterior-posterior directions is small[7], the apical area profile $A(n,t)$ is then equal to $L(n,t)$ multiplied by $A_0/L_0$, where $A_0 = 40\ \mu\text{m}^2$ is the cell apical area before gastrulation[6].

### Obtaining model parameter values by fitting model predictions to experimental measurements

To extract the damping length $\xi$ from experiment, we use as input to the model the experimentally measured myosin envelope from ref.[1], and compare the predicted apical area profile to the measured apical area profile from the same ref.[1]. To fit the characteristic contraction rate $w^*$, we used the experimentally measured myosin envelope at ~ 300 sec from ref.[2], and compared the predicted contraction rate profile (using the best fit $\xi$ from the previous fitting procedure) to the measured contraction rate profile at the same time from the same ref.[2].

The reason we used the data from ref.[1] to fit for the damping length $\xi$ is that the myosin levels and apical areas were measured for cells 1-10 in ref.[1], whereas in ref.[2], measurements were made for cells 1-7 only. Since $\xi$ is related to a length scale, it is more accurate to use measurements that include more cells. On the other hand ref.[2] measured myosin levels and apical areas at many instants in time, so these time-dependent apical area measurements can be used to estimate the instantaneous contraction rate profiles, and to fit for the characteristic contraction rate $w^*$ that sets the time scale of our system.

Before fitting, we first smoothed the experimental myosin envelopes from refs.[1,2] by non-linear least-squares fitting to a generic parameterized bell-shaped envelope (see Fig. S1A for the two fitted envelopes),

$$T_{\text{myo}}^{\text{env}}(n) = f^{an^4+bn^3+cn^2+dn+e}\,.$$

This curve fitted from ref.[1] is used across this study, referred to as the myosin envelope. For Fig. 2A, the myosin envelopes are assumed fixed in time. For Fig. 2B, we consider the realistic ramping up myosin envelope as model input, whose amplitude increases linearly in time from 0 sec, and the amplitude at $t = 300$ s is equal to 1 (this fitted curve above). For Figs. 3-4, we multiply this ramping up myosin envelope by stochastic factors $\big(1 + \epsilon_{\text{fluc}}(n,t)\big)$ which will be described below.

The myosin envelope for modeling *Spn27A* RNAi in Fig. 2A is a soft top-hat curve $T_{\text{myo}}^{\text{Spn27}A}(n) = \exp\left(-\left(\frac{n}{7}\right)^{10} / \ln 2\right)$ which mimics the experiment[2].

Next, we used the myosin envelope fitted from ref.[1] as input to the model and numerically solved the model, Eq. (2), to obtain the apical area profile (taking the myosin envelope to be fixed in time). We then performed a two-parameter least-squares fitting to extract the values of $\xi$ and $w^*$, by comparing this predicted apical area profile to the experimentally measured apical area profile from ref.[1]. This yielded a best fit $\xi = 2.5 \pm 1.2$ (mean ± standard error).

For the characteristic contraction rate $w^*$, we used the myosin profile measured at ~ 300 sec from ref.[2] and performed a one-parameter least-squares fitting to extract $w^*$ by comparing the predicted contraction rate profile to the measured contraction rate profile at the same time from ref.[2]. This yielded a best fit value $w^* = 42 \pm 6 \text{ nm s}^{-1}$. Using the experimentally measured apical myosin stress $T_{\text{myo}}^* = 1.5 \text{ nN}$ during dorsal closure[8], we estimated the internal viscosity $\mu = T_{\text{myo}}^*/w^* = 36 \pm 5 \text{ pN s nm}^{-1}$. We then estimated the external drag coefficient $\lambda = \mu/\xi^2 = T_{\text{myo}}^*/(w^*\xi^2) = 6.3 \pm 6.1 \text{ pN s nm}^{-1}$ (mean ± standard error).

#### Analytical results for three hypothetical myosin profiles

Here we present the analytical expressions for the contraction rate profiles $w(n)$ generated by the three hypothetical myosin profiles of Fig. 1 (top-hat, triangular and parabolic). The model Eq. (2) is written

$$w''(n) - \frac{1}{\xi^2} w'(n) = w^* T_{\text{myo}}''(n).$$

We solve the above equation using the Green's function obtained from the equation $G''(n) - G(n)/\xi^2 = \delta(n)$. The Green's function reads

$$G(n) = -\frac{\xi}{2} e^{-\frac{|n|}{\xi}}.$$

Given a myosin profile $T_{\text{myo}}(n)$, the analytical solution of the contraction can be obtained by the convolution $G(n) * \left(w^* T_{\text{myo}}''(n)\right)$.

##### Top-hat myosin profile

For the top-hat myosin profile $T_{\text{myo}}(n) = \Theta(n + h/2) - \Theta(n - h/2)$ (Fig. 1E, left). Therefore, we solve the contraction rate profile,

$$w(n) = \frac{w^*}{2}\left(\frac{n+h/2}{|n+h/2|}e^{-\frac{|n+h/2|}{\xi}} - \frac{n-h/2}{|n-h/2|}e^{-\frac{|n-h/2|}{\xi}}\right).$$

**Triangular myosin profile (constant magnitude gradient)**

For a triangular myosin profile (Fig. 1E, middle), we have myosin profile

$$T_{\mathrm{myo}}(n) = \begin{cases} 1-|n/h|, & |n| \le h \\ 0, & |n| > h \end{cases}.$$

This gives the contraction rate profile

$$w(n) = \frac{w^*\xi}{2h}\left(2e^{-\frac{|n|}{\xi}} - e^{-\frac{|n+h|}{\xi}} - e^{-\frac{|n-h|}{\xi}}\right).$$

**Parabolic myosin profile (non-zero curvature)**

For the parabolic myosin profile (Fig. 1E, right), we have

$$T_{\mathrm{myo}}(n) = \begin{cases} -2n^2/h^2 + 1, & |n| \le h/\sqrt{2} \\ 0, & |n| > h/\sqrt{2} \end{cases}.$$

We solve the contraction rate profile (for $n \ge 0$)

$$w(n) = -w^*\frac{\sqrt{2}\xi}{h}\left(e^{-\frac{\left|n-\frac{h}{\sqrt{2}}\right|}{\xi}} + e^{-\frac{\left|n+\frac{h}{\sqrt{2}}\right|}{\xi}}\right) + \begin{cases} 2w^*\left(\frac{h}{\xi}\right)^2\left(2 - e^{-\frac{\frac{h}{\sqrt{2}}-n}{\xi}} - e^{-\frac{\frac{h}{\sqrt{2}}+n}{\xi}}\right), & 0 \le n < \frac{h}{\sqrt{2}} \\ 2w^*\left(\frac{h}{\xi}\right)^2\left(e^{\frac{h}{\sqrt{2}\xi}} - e^{-\frac{h}{\sqrt{2}\xi}}\right)e^{-\frac{n}{\xi}}, & n \ge \frac{h}{\sqrt{2}} \end{cases}$$

The full contraction rate profile is the above expression together with its even extension into $n < 0$.

#### Extracting the amplitude of fluctuations in cell apical myosin levels from experiment

Fluctuations in cell myosin levels are in important feature of our model. Their magnitude is governed by the parameter $\epsilon_{\mathrm{fluc}}$, which represents the size of the fluctuations relative to the mean myosin value. Here we summarize the procedure we used to estimate the value of $\epsilon_{\mathrm{fluc}}$ from experimental measurements. From the reported experimental myosin profile of ref.[1] (see Fig. S6A), we first extracted the median myosin level for each cell number $n$. We then assumed that for each cell number $n$ the myosin level follows a normal distribution with mean equal to the median. Then, from the box size ($25^{\mathrm{th}}$-$75^{\mathrm{th}}$ percentiles) for each cell in Fig. S3A, we estimated the standard deviations of myosin levels for each cell number $n$ (Fig. S6B). We assumed the relative fluctuation amplitude $\epsilon_{\mathrm{fluc}}$ (standard deviation divided by the mean) is the same for all cells, and found a best-fit value $\epsilon_{\mathrm{fluc}} = 46 \pm 8\%$ (mean ± standard error) by least-square fitting (Fig. S6B). Thus, as a value representative of this experimental data, in our model we used a value $\epsilon_{\mathrm{fluc}} = 45\%$.

### The stochastic factor $\epsilon_{\mathrm{fluc}}$ to represent myosin signals

To reproduce the myosin signals in wild-type and *C-GAP* RNAi embryos, the spatiotemporal myosin profile $T_{\mathrm{myo}}(n,t)$ is modeled as

$$T_{\mathrm{myo}}(n,t) = \big(1 + \epsilon_{\mathrm{fluc}}(n,t)\big) T_{\mathrm{myo}}^{\mathrm{env}}(n,t).$$

where $T_{\mathrm{myo}}^{\mathrm{env}}(n,t)$ is the ramping up myosin envelope with no fluctuations. This envelope profile is multiplied by a stochastic factor $1 + \epsilon_{\mathrm{fluc}}(n,t)$, where $\epsilon_{\mathrm{fluc}}(n,t)$ is a random variable following a normal distribution, uncorrelated from cell to cell ($n$). The time correlation has the form

$$\langle \epsilon_{\mathrm{fluc}}(n,t)\, \epsilon_{\mathrm{fluc}}(n',t+\Delta t)\rangle_c = \langle {\epsilon_{\mathrm{fluc}}}^2\rangle_c\, \delta_{n,n'} \exp\left(-\frac{\Delta t}{\tau}\right).$$

In other words, for each cell $n$, the stochastic factor $\epsilon_{\mathrm{fluc}}(n,t)$ is the trajectory of a Brownian particle within a harmonic potential – the Ornstein-Uhlenbeck process. The amplitude of the fluctuations is taken to be $\langle \epsilon_{\mathrm{fluc}}^2 \rangle_c^{1/2} = 0.45$ for both wild-type and *C-GAP* RNAi embryos, estimated from experiment (see section above, *Extracting the amplitude of fluctuations in cell apical myosin levels from experiment*). For wild-type, we took the correlation timescale to be $\tau = 20$ s, so that a typical pulse has duration ~ 1 min, consistent with experiment[9]. For *C-GAP* RNAi embryos, we took $\tau = 300$ s to mimic the experimental[10] non-pulsatile myosin dynamics (see Fig. 3B). See Fig. S6C for two realizations of the stochastic factor $\epsilon_{\mathrm{fluc}}(n,t)$ versus time for wild-type.
